# Illness, social disadvantage, and sexual risk behavior in adolescence and the transition to adulthood

**DOI:** 10.1101/419010

**Authors:** Jenna Alley, Rebecca Y. Owen, Sarah E. Wawrzynski, Lauren Lasrich, Zobayer Ahmmad, Rebecca Utz, Daniel E. Adkins

## Abstract

This study investigated the influence of illness on sexual risk behavior in adolescence and the transition to adulthood, both directly and through moderation of the impact of social disadvantage. We hypothesized positive effects for social disadvantages and illness on sexual risk behavior, consistent with the development of faster life history strategies among young people facing greater life adversity. Using the first two waves of the National Longitudinal Study of Adolescent Health, we developed a mixed effects multinomial logistic regression model predicting sexual risk behavior in three comparisons, risky nonmonogamous sex vs. 1) safer nonmonogamous sex, 2) monogamous sex, and 3) abstinence, by social characteristics, illness, interactions thereof, and control covariates. Multiple imputation was used to address a modest amount of missing data. Subjects reporting higher levels of illness had lower odds of having safer nonmonogamous sex (OR = 0.84, *p* < .001), monogamous sex (OR = 0.82, *p* < .001), and abstinence (OR = 0.74, *p* < .001) vs. risky nonmonogamous sex, relative to individuals in better health. Illness significantly moderated the sex (OR = 0.88, *p* < .01), race/ethnicity (e.g., OR = 1.21, *p* < .001), and childhood SES (OR = 0.94; *p* < .01) effects for the abstinent vs. risky nonmonogamous sex comparison. Substantive findings were generally robust across waves and in various sensitivity analyses. These findings offer general support for the predictions of life history theory. Illness and various social disadvantages are associated with increased sexual risk behavior in adolescence and the transition to adulthood. Analyses indicate that the buffering effects of several protective social statuses against sexual risk-taking are substantially eroded by illness.

## INTRODUCTION

Understanding the determinants of sexual risk behavior has important public health implications given associations between sexual risk-taking and negative health and quality of life outcomes, including STD/STI and unintended pregnancy (Edwards & Coleman, 2004; Ryan, Mendle, & Markowitz, 2015). Previous research has shown childhood adversity to be strongly associated with increased risk behavior during adolescence and young adulthood, including risky sexual behavior (Simpson, Griskevicius, Kuo, Sung, & Collins, 2012; Del Giudice, Gangestad, & Kaplan, 2015). Adolescence is of particular interest in the study of sexual behavior as it is the cusp of sexuality and fertility, characterized by critical transitions including puberty, sexual debut, and sexual identity formation (Coley, Votruba Drzal, & Schindler, 2009). Relatedly, adolescence is characterized by a normative increase in risk behavior (Coley et al., 2009), particularly among males and various socially disadvantaged groups (Hagan & Foster, 2003). Life history theory (LHT) offers a compelling paradigm for investigating this process, as it provides a comprehensive theoretical explanation with testable predictions, regarding the influence of adversity on sexual risk behavior in adolescence and young adulthood.

LHT posits that individuals make biobehavioral trade-offs in their allocation of time and energy to maximize the primary evolutionary outcomes of survival and reproduction (i.e., fitness) (Hill, 1993). These trade-offs manifest as developmental adaptions to the environment, which promote fitness via behavioral strategies calibrated by early social experiences (Chisholm, 1999; Chisholm et al., 1993). Specifically, as early social adversity predicts later extrinsic mortality, the theory posits that individuals are calibrated by early social cues to accelerate/decelerate life history trait development to match their predicted extrinsic mortality risk (Chisholm et al., 1993; Belsky, Steinberg, & Draper 1991). A primary manifestation of this calibration during adolescence and young adulthood is theorized to be the trade-off of mating vs. parenting effort, indicated by behavioral traits including earlier sexual debut and greater number of sexual partners (Ellis, McFadyen-Ketchum, Dodge, Pettit, & Bates, 1999; Griskevicius, Delton, Robertson, & Tybur, 2011). Thus, LHT posits that experiencing more adverse early environments predisposes individuals to maximize short-term mating, achieved during adolescence by earlier sexual debut and increased sociosexuality (i.e., willingness to have casual sex, and to engage in sex without love, closeness, or commitment) (Szepsenwol et al., 2017). A substantial body of empirical research supports this perspective; for instance, indicating that early life stress accelerates life history speed, in part by fostering unrestricted sociosexual attitudes (e.g., Patch & Figueredo, 2017; Szepsenwol et al., 2017).

The sum of an individual’s life history trade-offs across development may be conceptualized as their life history strategy (e.g., Del Giudice et al., 2015; Ellis, Figueredo, Brumbach, & Schlomer, 2009). Life history strategies are typically conceptualized as a continuum of “fast” to “slow”, calibrated by one’s extrinsic mortality risk, as indicated by the level of unpredictability and/or harshness in one’s (primarily early) environment (Kaplan & Gangestad, 2005). Previous research has offered support for this perspective, including association of faster life histories to unpredictable environments, including parental job loss, divorce/conflict, and frequent changes in household composition, as well as harsh environmental exposures, such as childhood maltreatment, poverty, and racial discrimination (Belsky, Schlomer, & Ellis, 2012; Schafer, Ferraro, & Mustillo, 2011; Simpson et al., 2012). Thus, LHT theorizes that early experiences of social disadvantage signal increased mortality risk, which encourages faster biobehavioral development of life history traits (Ellis, Oldehinkel, & Nederhof, 2017; Szepsenwol et al., 2017).

Recent research has argued that the signal of increased extrinsic mortality risk need not be restricted to external social cues. Rather, a growing literature within LHT has proposed that internal states of health and illness can also signal increased mortality risk and, thus, trigger developmental shifts toward fast life history strategies (Griskevicius et al., 2011). Proposed indicators of such adverse internal states include chronic illness, functional limitations, and disability (Griskevicius et al., 2011; Waynforth, 2012). Thus, it is posited that an internal predictor, such as chronic and/or severe physical illness, can signal increased mortality risk, which fosters biobehavioral adaptation toward fast life history strategies, including maximizing short-term mating at the expense of longer-term parental investment (Hill, Boehm, & Prokosch, 2016; Waynforth, 2012). Arnocky, Pearson, & Vaillancourt (2015) provide compelling support for this internal prediction model, showing that an individual’s history of illness can influence jealousy and perceptions of fidelity, important psychological correlates of fast life history sexual strategies. Specifically, they found that individuals with more severe health problems reported more jealousy towards their partner and higher perceptions of potential infidelity on the part of their partner. These findings suggest that internal states of health and illness may have a multifaceted influence on mating behaviors, influencing personal sexual decisions (Griskevicius et al., 2011), as well as mate retention strategies (Arnocky, Pearson, & Vaillancourt, 2015).

While the main effect of respondent sex, and interactive effect of sex and illness, are not a primary focus of this study, given the important role of sex and gender in shaping sexual behavior, these processes are considered. Some sex differences in sexual behavior are well-established, including the observation that males generally exhibit greater interest in, and receptivity to, casual sex than their female counterparts (e.g., Clark & Hatfield, 1989), likely due, in part, to sex differences in parental investment (Trivers, 1972). Previous research has also shown that early environmental conditions of adversity appear to have a particularly strong influence on female’s reproductive strategies, compared to males (e.g., Fergusson, Horwood, & Lynskey, 1997; Haydon, Hussey, & Halpern, 2011; James, Ellis, Schlomer, & Garber, 2012). While various theoretical explanations of this phenomena have been advanced, the leading LHT perspective argues that females exhibit greater sexual behavioral plasticity in response to adversity because females’ optimal fitness maximizing strategy is more environmentally contingent than males (or was in the evolutionary environment) (Ellis et al., 2009). That is, the availability of social and financial resources from partner and family is disproportionately important to female childbearing and rearing outcomes relative to males (or was in the evolutionary environment), due, in part, to sex differences in baseline parental investment. Consistent with this theoretical perspective, empirical work has shown that the experience of adversity is associated with increased female perceptions of sexual intent and increased interest in casual sex (DelPriore, Proffitt Leyva, Ellis, & Hill, 2018).

### Hypotheses

The current study aims to elaborate the internal predictors perspective within LHT. To this end, we hypothesize: H1: Individuals who experience more social disadvantage (e.g., childhood trauma, low socioeconomic status, minority stress) will be more likely to engage in risky nonmonogamous sex than those who experience greater social advantage. H2: Illness will be positively associated with engaging in risky nonmonogamous sex. H3: Illness will attenuate the buffering effects of social advantage on sexual risk-taking.

## METHOD

### Participants

We used data from the first two waves of the National Longitudinal Study of Adolescent Health (Add Health). Add Health is the largest longitudinal sample of contemporary adolescents to young adults in the United States. Sample features have been previously described (http://www.cpc.unc.edu/projects/addhealth/design). Briefly, a sample of 20,745 adolescents in grades 7–12 were initially assayed for data collection in 1994–1995 (Wave 1), and then again 18 months later in 1996 (Wave 2). The age range across the two waves was approximately 11.6-22.5, with a mean of 16.7. In Wave 1, a questionnaire was administered to a selected residential parent of each adolescent, from which some covariates in the present analysis were derived (e.g., childhood household income, parental educational attainment). The use of multiple imputation (see below) necessitated the analysis of unweighted data, an analytical decision supported by current statistical thinking on the use of weights in survey data (Bollen, Biemer, Karr, Tueller, & Berzofsky, 2016). Add Health participants provided written, informed consent for participation in all aspects of Add Health per the University of North Carolina School of Public Health Institutional Review Board guidelines (https://www.cpc.unc.edu/projects/addhealth/faqs/index.html#Was-informed-consent-required). This analysis has been approved by the University of Utah Institutional Review Board (IRB_00107767).

### Measures

#### Sexual risk behavior

We used several items from Add Health, measured at each of the two analyzed waves, to classify subjects’ sexual behavior as one of four exhaustive and mutually exclusive categories: risky nonmonogamous sex, safer nonmonogamous sex, monogamous sex, and abstinence. The first item used asked whether the individual has ever had sex; individuals responding no to this item were coded as abstinent. The second item asked if the individual has ever had sex outside of a romantic relationship in the past 12 months. Individuals reporting yes to the first item and no to the second items were coded as “monogamous sex”. We then used an item asking if the subject used contraception in most sexual encounters, and a pair of questions asking if the subject used a condom in the first and most recent sexual encounter to parse the nonmonogamous group into “risky” and “safer” categories. Specifically, if a subject reported having sex outside of a romantic relationship and either regular contraceptive use or regular condom use (i.e., using a condom in both their first and most recent sexual encounter), they were coded as safer nonmonogamous. Conversely, individuals reporting having sex outside of a relationship and inconsistent (or non-) contraceptive use were classified as risky nonmonogamous sex. Logical inconsistencies between items were coded missing and addressed by multiple imputation (see below).

While some previous LHT-relevant sexuality research has examined sociosexuality using scales such as the Sociosexual Orientation Inventory (Penke & Asendorpf, 2008), Add Health does not contain a specific sociosexuality instrument. And though our measure of sexual risk taps a central aspect of sociosexuality—sexual activity outside of a committed relationship (Penke & Asendorpf, 2008), our measure is not at all synonymous with the sociosexuality construct. Rather, our study has a primary focus on sexual risk (to health and well-being) that only partially overlaps with sociosexual behavior. This is because sociosexual behavior, when paired with consistent contraceptive use, is no riskier than monogamous sexual behavior. So, while we theorize sociosexuality as a mechanism in the adversity sexual risk process modeled, we are primarily concerned with the more distal outcome of sexual risk behavior.

#### Illness

Illness was assessed in both waves using a 5-level self-rated health item. Specifically, subjects were asked to respond to the question, “In general, how would you rate your health?” Response options ranged from “Poor” to “Excellent.” Previous research has consistently, and somewhat surprisingly, demonstrated that this simple operationalizion of self-rated health/illness is strongly predictive of future mortality and morbidity outcomes (Benyamini & Idler, 1999; Idler & Benyamini, 1997). Previous analyses of the current data (i.e., Add Health) have demonstrated that this measure of self-rated health is strongly associated with a range of physical health measures (Boardman, 2006). The measure was standardized in analyses.

#### Severe childhood adversity

Measures of childhood adversity comprised five dichotomous indicators: sexual abuse, physical abuse, neglect, drugs/alcohol accessible at home, and parental incarceration, occurring in adolescence. These five severe childhood adversities are described in detail in Supplemental Methods. Briefly, each of the five childhood adversity measures retrospectively assessed childhood and adolescent experiences using multiple questionnaire items. A cumulative adversity count ranging from 0 to 5 was used in the primary analysis; sensitivity analyses examining the effects of the individual childhood adversities are presented in Supplemental Tables S6a-c.

#### Sociodemographic characteristics

All presented models included a set of sociodemographic covariates characterizing the adolescent’s social background. These included household SES measures for parental education and household income assessed at Wave 1. These measures have been previously described (Adkins, Wang, Dupre, van den Oord, & Elder, 2009), and are described in detail in Supplemental Methods. Both parental education and income were standardized in the analysis. Self-reported race/ethnicity (coded Black, Asian, Hispanic, White (ref), and Other), nativity (i.e., first-generation immigrant status), and age were also included as covariates in all presented models. Sex was self-reported by the Add Health subjects, as either male or female, in Wave 1. Finally, sexual attraction was constructed as a four category nominal variable (i.e., asexual, bisexual, same-sex only, and opposite-sex only attraction) using two questions asking the subject whether they felt attracted to a) males and b) females, in combination with the subject’s sex.

### Statistical Analyses

There was a small to moderate amount of missing data at both waves (< 30% missing for all analysis variables and < 4% for most; see Supplemental Table S2). We addressed missing data using the multiple imputation (MI) method, multiple imputation with chained equations (MICE) (White, Royston, & Wood, 2011). We used a conservative 30 imputations in all MI analyses (von Hippel, 2018). Given the large size of the Add Health data and the relatively modest missingness of the current analytical sample, the primary advantage of MI was not maximizing efficiency (and, thus, statistical power), but instead minimizing the most likely sources of parameter estimate bias. That is to say, MI has optimal properties in both the case of data “missing at random” (MAR) and “missing completely at random” (MCAR), and generally performs comparably to listwise deletion in scenarios of “not missing at random” (NMAR) (Little & Rubin, 2019; Rubin, 2004). Many of the variables included in our model (e.g., illness, race/ethnicity, sexual orientation, SES) are associated with missingness, so the data are certainly MAR to some, nontrivial degree. And while the degree to which the data are NMAR is inherently empirically untestable, analytical and simulation studies suggest that MI has optimal properties relative to listwise deletion in the majority of likely scenarios (Schafer & Graham, 2002; van Ginkel, Linting, Rippe, & van der Voort, 2019). Additional MI methodological details are presented in Supplemental Methods.

Our primary analytical approach pooled data from Waves 1 and 2 to maximize statistical power (wave-specific sensitivity analyses are also reported in Supplemental Tables S3a-c and S4a-c). Thus, primary inferential analyses used mixed effects multinomial logistic regression, which was used to model a polytomous outcome variable with 4 categories, yielding 3 logistic comparisons (risky nonmonogamous sex vs. safer nonmonogamous sex, monogamous sex, and abstinence) using the common reference category of risky nonmonogamous sex. In each of these comparisons, the logged odds of the outcomes were predicted by a linear combination of the predictor variables, with a subject-level random intercept to account for the multilevel data structure of observations nested within subjects, which corrects standard errors and coefficients for nonindependence of observations (Singer & Willett, 2003). This approach can essentially be conceptualized as a series of logistic regressions predicting sexual behavior outcomes, versus a common reference category (i.e., risky nonmonogamous sex), by social characteristics, illness, interactions thereof, and controls.

The model was developed beginning with (1) a baseline model of social disadvantage and sociodemographic controls predicting the (logged odds of) four category sexual risk outcome. We followed this baseline model with (2) a model adding illness as a predictor. Next, in order to test whether illness moderates the effects of the social factors, we (3) add interactions between illness and all of the significantly associated social factors from Model 2. In the final model, (4) we trim nonsignificant (p ≥.01) interactions from the model. The full model is visualized in Figure 1.

**Figure 1.**
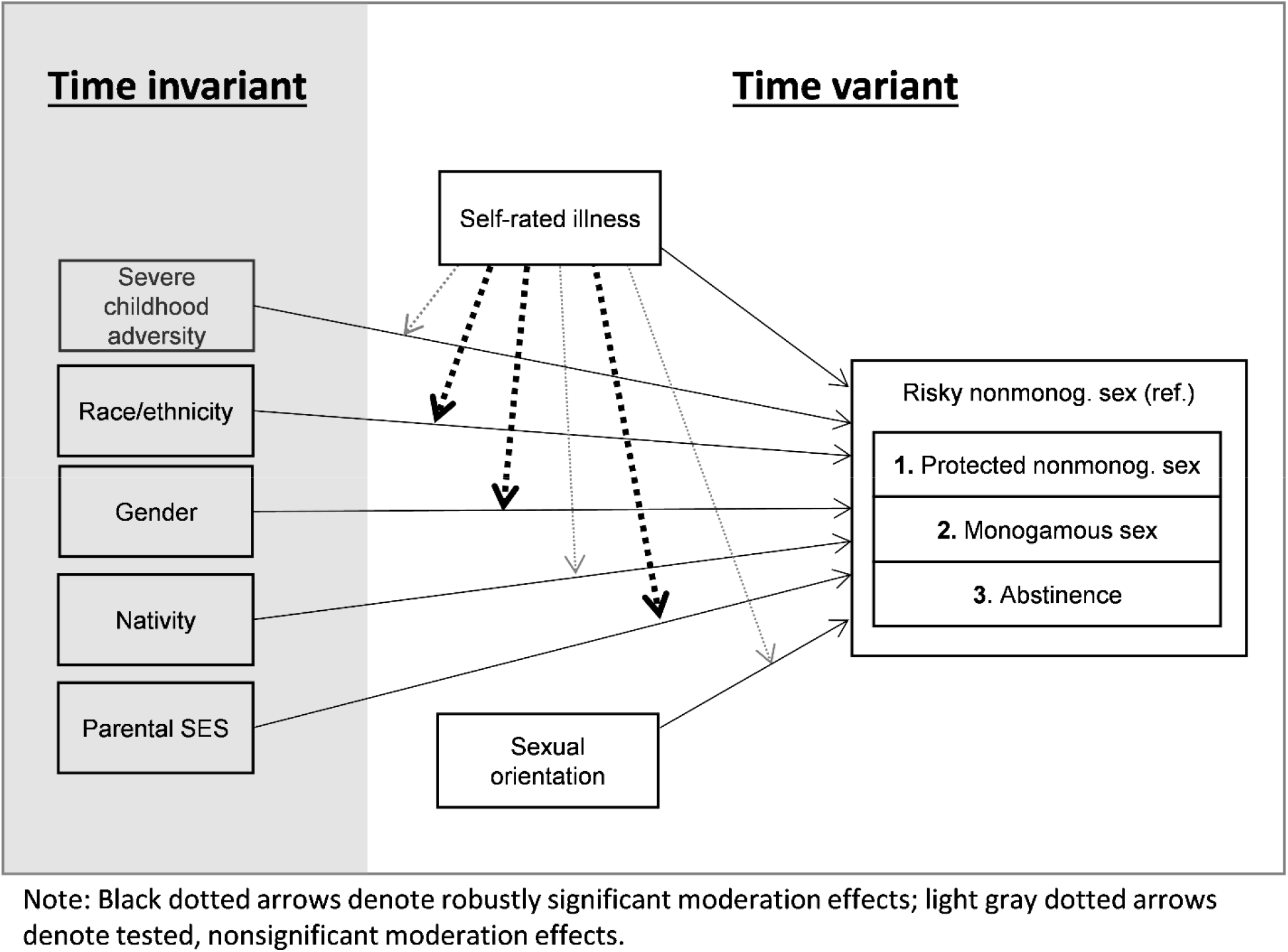
Simplified path diagram of the proposed model of moderation, by physical illness, of the effects of social disadvantage on sexual risk behavior. Bold dotted lines represent statistically significant moderation effects, thin dotted lines represent nonsignificant tested moderation effects

To test the robustness of our substantive results, we conducted several sensitivity analyses. These are briefly discussed in the results and documented in Supplementary Materials. Sensitivity analyses include the following: (A) the primary model sequence using complete case analysis (rather than multiple imputation). (B) The primary model sequence stratified by wave. (C) A model substituting the five individual severe childhood adversities (i.e., sexual abuse, physical abuse, neglect, drugs/alcohol in the home, parental incarceration) for the cumulative childhood adversity index. To support research rigor and reproducibility, the data management and analysis script are available through the Open Science Framework (???).

## RESULTS

Descriptive statistics for the analysis variables are detailed in Table 1. Briefly, abstinent was the modal sexual behavior category in both waves I (60%) and II (51%). The monogamous sexual behavior category increased in frequency, notably across waves from 17% (WI) to 32% (WII). Both risky (WI: 11%; WII: 8%) and safer nonmonogamous (WI: 12%; WII: 9%) categories became less frequent. Illness was standardized in both waves, as was parental education and logged household income. Demographic proportions are approximately reflective of national proportions at the time. The age range across the two waves was approximately 11.6-22.5, with a mean of 16.7. The mean number of severe childhood adversities was 0.7 and ranged 0-5. Opposite sex-only attraction was the modal sexual attraction category and decreased in prevalence from WI (82%) to WII (75%). Reflecting the developmental period, asexual attraction ranged 12-20%, while bisexual attraction ranged 5-3%, and same-sex only attraction was steady at ∼1% in both waves.

**Table 1.** Descriptive statistics of analysis variables (MI, m = 30; N_j_ = 20774)

| Variable | Mean | SD | min | max |
| --- | --- | --- | --- | --- |
| Wave I: Risky nonmonogamous sex | 0.11 | 0.31 | 0 | 1 |
| Wave II: Risky nonmonogamous sex | 0.08 | 0.27 | 0 | 1 |
| Wave I: Safer nonmonogamous sex | 0.12 | 0.32 | 0 | 1 |
| Wave II: Safer nonmonogamous sex | 0.09 | 0.28 | 0 | 1 |
| Wave I: Monogamous sex | 0.17 | 0.38 | 0 | 1 |
| Wave II: Monogamous sex | 0.32 | 0.47 | 0 | 1 |
| Wave I: Abstinent | 0.60 | 0.49 | 0 | 1 |
| Wave II: Abstinent | 0.51 | 0.50 | 0 | 1 |
| Wave I: Illness | -0.02 | 1.00 | -2.19 | 3.17 |
| Wave II: Illness | -0.04 | 0.99 | -3.55 | 3.80 |
| Wave I: Asexual attraction | 0.12 | 0.32 | 0 | 1 |
| Wave II: Asexual attraction | 0.20 | 0.40 | 0 | 1 |
| Wave I: Bisexual attraction | 0.05 | 0.22 | 0 | 1 |
| Wave II: Bisexual attraction | 0.03 | 0.18 | 0 | 1 |
| Wave I: Opposite sex only attraction | 0.82 | 0.39 | 0 | 1 |
| Wave II: Opposite sex only attraction | 0.75 | 0.43 | 0 | 1 |
| Wave I: Same sex only attraction | 0.01 | 0.11 | 0 | 1 |
| Wave II: Same sex only attraction | 0.01 | 0.12 | 0 | 1 |
| Wave I: Age | 16.15 | 1.74 | 11.66 | 21.00 |
| Wave II: Age | 17.30 | 1.82 | 12.63 | 22.50 |
| Immigrant | 0.08 | 0.27 | 0 | 1 |
| Hispanic | 0.17 | 0.38 | 0 | 1 |
| Black | 0.22 | 0.42 | 0 | 1 |
| Asian | 0.07 | 0.26 | 0 | 1 |
| Other Race | 0.03 | 0.17 | 0 | 1 |
| White | 0.50 | 0.50 | 0 | 1 |
| Female | 0.51 | 0.50 | 0 | 1 |
| Household Income | -0.02 | 1.00 | -4.29 | 4.15 |
| Parental Education | 0.00 | 1.00 | -3.53 | 2.86 |
| Cumulative Severe Adversity | 0.70 | 0.95 | 0 | 5 |

Primary inferential analyses comprised a series of four nested, mixed effects multinomial logistic regressions comparing the odds of engaging in safer nonmonogamous sex (Comparison 1), monogamous sex (Comparison 2), and abstinence (Comparison 3), versus the reference category—risky nonmonogamous sex. Given the large number of parameters estimated by these multinomial logistic regressions, we primarily focus discussion on the coefficients directly relevant to our hypotheses and avoid discussion of most nonsignificant coefficients.

### Model 1: Social Disadvantage

The baseline model in the series includes sociodemographic characteristics (e.g., nativity, race/ethnicity, sex, childhood SES), sexual attraction, and cumulative severe childhood adversity, in order to map the effects of social disadvantage on sexual risk behavior. Regarding the safer vs. risky nonmonogamous comparison shown in Table 2, Model 1, subjects identifying as Hispanic (OR = 0.78, *p* < .01), Black (OR = 0.86, *p* < .05), and Asian (OR = 0.51, *p* < .001) exhibited lower probabilities of engaging in safer vs. risky nonmonogamous sex, compared to Whites. Females exhibited marginally lower probability of engaging in safer vs. risky nonmonogamous sex, though this was not robust across the other model specifications in the series. The two childhood SES indicators were both associated with increased probability of safer (vs. risky) nonmonogamous sex (income: OR = 1.10, *p* < .01; parental education: OR = 1.11, *p* < .01). The severe adversity index was strongly associated with risky nonmonogamous sex (OR = 0.87, *p* < .001), in this comparison.

**Table 2.**
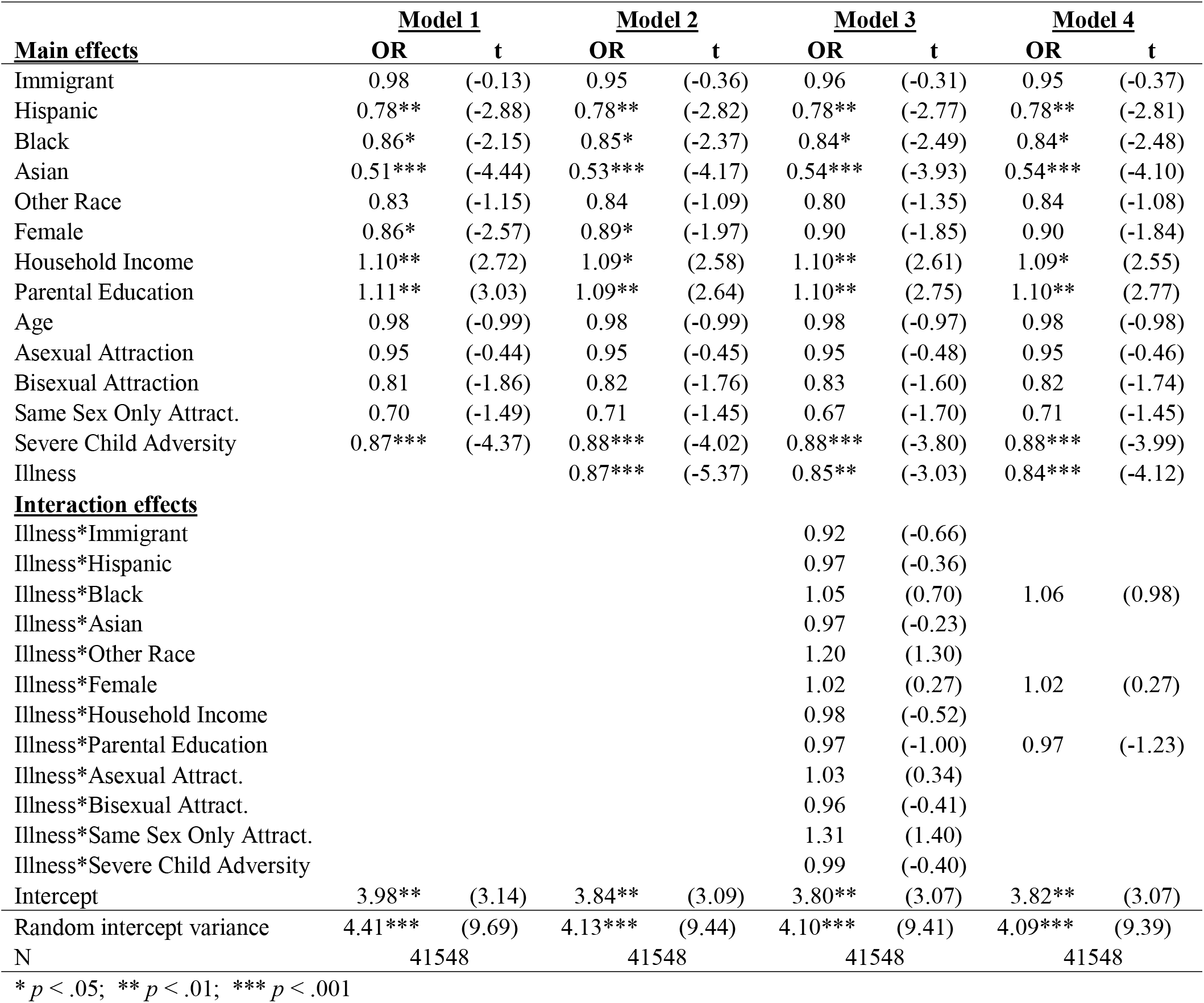
Multinomial logistic regression of pooled wave I & wave II data: Protected nonmonogamous sex v. risky nonmonogamous sex (MI, m = 30)

In the monogamous vs. risky nonmonogamous comparison (Table 3, Model 1), the direction of effect was the same and the magnitude of effects was greater than in the safer vs. risky nonmonogamous comparison, with a few exceptions. Specifically, Black and “Other Race” (primarily multiracial) youth exhibited decreased likelihood of monogamous (vs. risky monogamous) sex, compared to Whites; and female sex was strongly significantly associated with monogamous sex vs. risky nonmonogamous sex. Age was positively associated with monogamous sex vs. risky nonmonogamous sex (OR = 1.07, *p* < .01). Asexual attraction was strongly associated with monogamous sex (OR = 2.45, *p* < .001), while bisexual attraction was associated with risky nonmonogamous sex (OR = 0.61, *p* < .001).

**Table 3.**
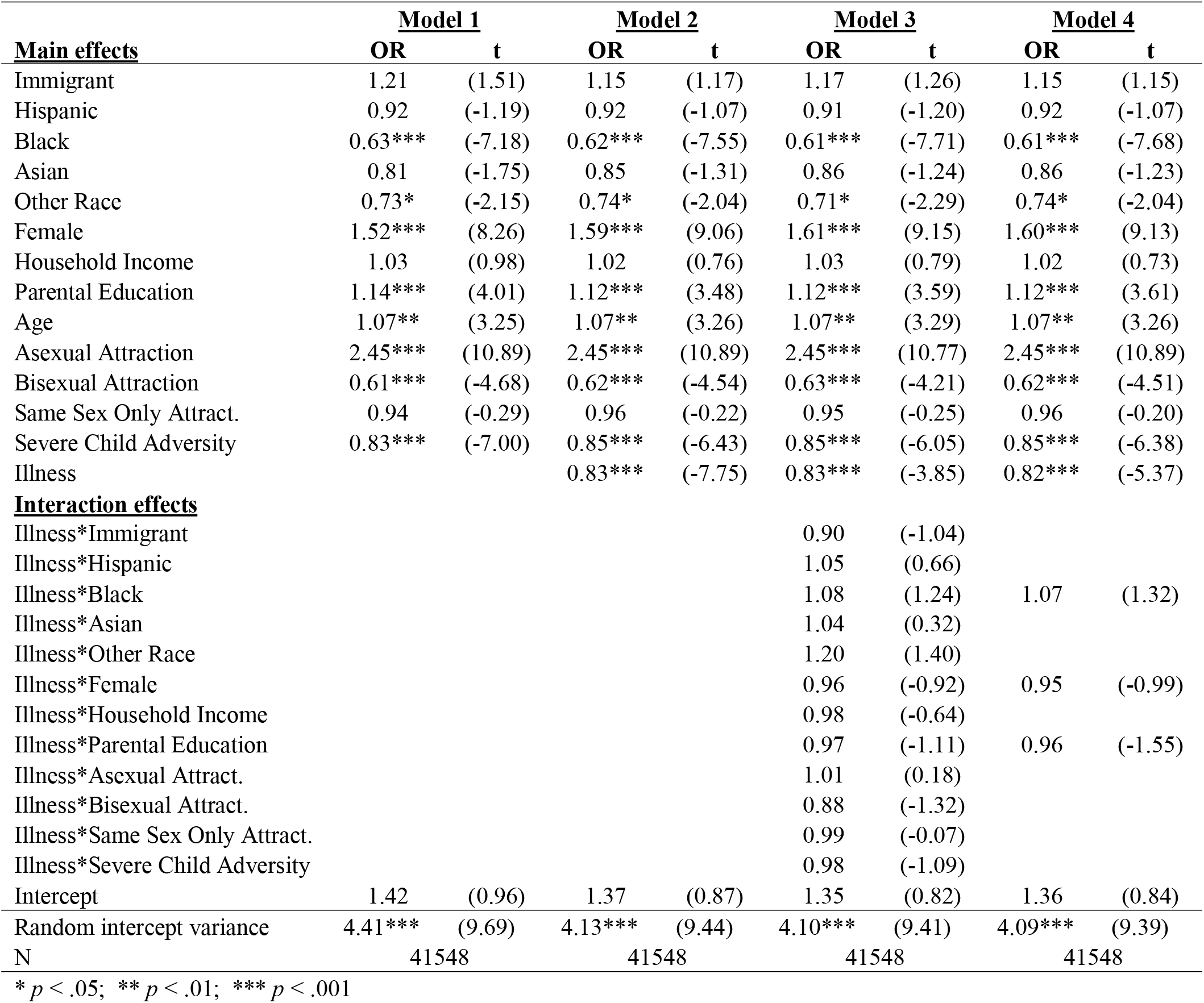
Multinomial logistic regression of pooled wave I & wave II data: Monogamous sex v. risky nonmonogamous sex (MI, m = 30)

In the abstinence vs. risky nonmonogamous sex comparison (Table 4, Model 1), the direction of effect was the same and the magnitude of effects was greater than in the monogamous vs. risky nonmonogamous comparison, with a single exception—identifying as a first-generation immigrant was strongly associated with abstinence vs. risky nonmonogamous sex (OR = 2.26, *p* < .001).

**Table 4.**
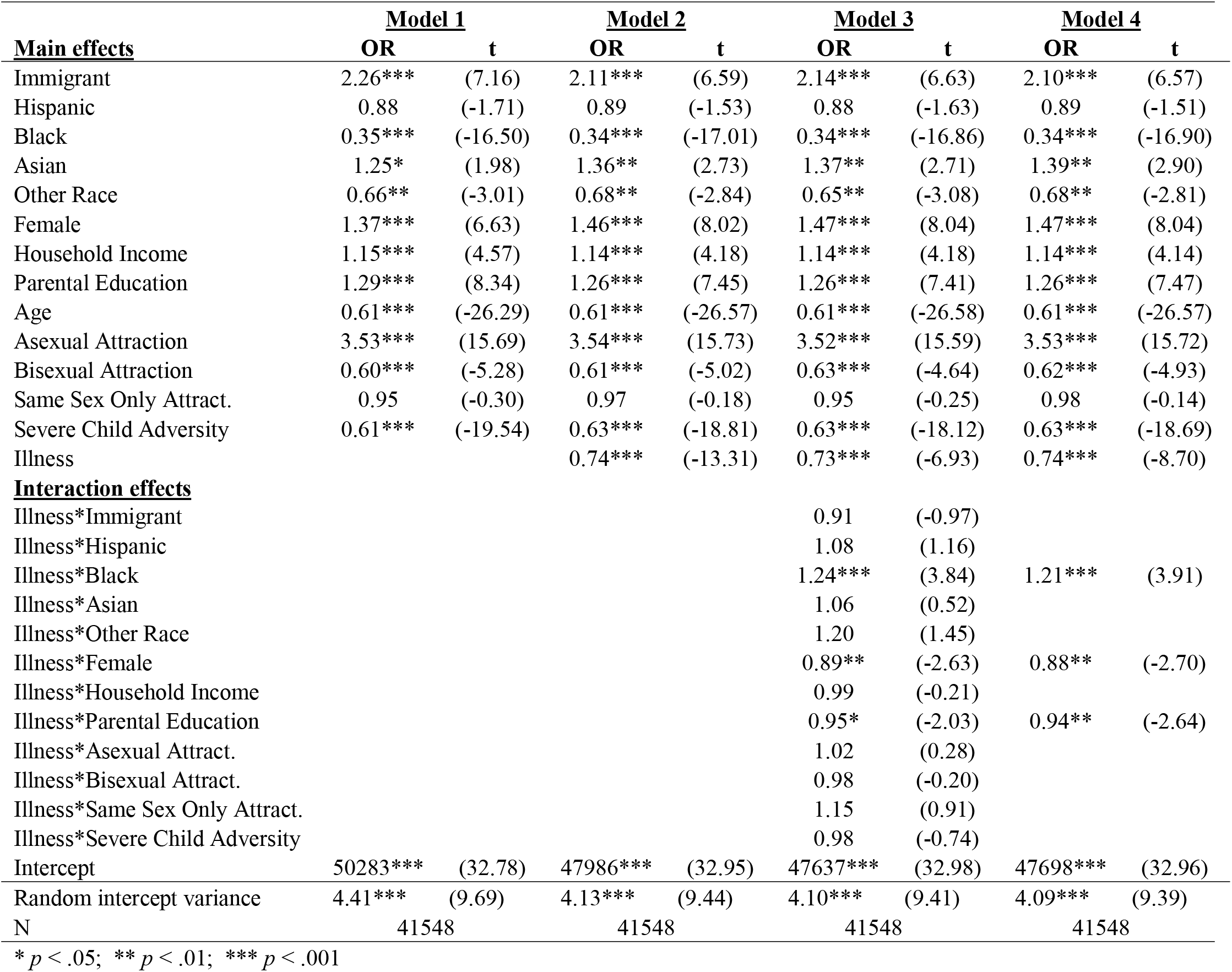
Multinomial logistic regression of pooled wave I & wave II data: Abstinence v. risky nonmonogamous sex (MI, m = 30)

|  | <u>Model 1</u> |  | <u>Model 2</u> |  | <u>Model 3</u> |  | <u>Model 4</u> |  |
| --- | --- | --- | --- | --- | --- | --- | --- | --- |
| <u>Main effects</u> | OR | t | OR | t | OR | t | OR | t |
| Immigrant | 2.26*** | (7.16) | 2.11*** | (6.59) | 2.14*** | (6.63) | 2.10*** | (6.57) |
| Hispanic | 0.88 | (-1.71) | 0.89 | (-1.53) | 0.88 | (-1.63) | 0.89 | (-1.51) |
| Black | 0.35*** | (-16.50) | 0.34*** | (-17.01) | 0.34*** | (-16.86) | 0.34*** | (-16.90) |
| Asian | 1.25* | (1.98) | 1.36** | (2.73) | 1.37** | (2.71) | 1.39** | (2.90) |
| Other Race | 0.66** | (-3.01) | 0.68** | (-2.84) | 0.65** | (-3.08) | 0.68** | (-2.81) |
| Female | 1.37*** | (6.63) | 1.46*** | (8.02) | 1.47*** | (8.04) | 1.47*** | (8.04) |
| Household Income | 1.15*** | (4.57) | 1.14*** | (4.18) | 1.14*** | (4.18) | 1.14*** | (4.14) |
| Parental Education | 1.29*** | (8.34) | 1.26*** | (7.45) | 1.26*** | (7.41) | 1.26*** | (7.47) |
| Age | 0.61*** | (-26.29) | 0.61*** | (-26.57) | 0.61*** | (-26.58) | 0.61*** | (-26.57) |
| Asexual Attraction | 3.53*** | (15.69) | 3.54*** | (15.73) | 3.52*** | (15.59) | 3.53*** | (15.72) |
| Bisexual Attraction | 0.60*** | (-5.28) | 0.61*** | (-5.02) | 0.63*** | (-4.64) | 0.62*** | (-4.93) |
| Same Sex Only Attract. | 0.95 | (-0.30) | 0.97 | (-0.18) | 0.95 | (-0.25) | 0.98 | (-0.14) |
| Severe Child Adversity | 0.61*** | (-19.54) | 0.63*** | (-18.81) | 0.63*** | (-18.12) | 0.63*** | (-18.69) |
| Illness |  |  | 0.74*** | (-13.31) | 0.73*** | (-6.93) | 0.74*** | (-8.70) |
| <u>Interaction effects</u> |  |  |  |  |  |  |  |  |
| Illness*Immigrant |  |  |  |  | 0.91 | (-0.97) |  |  |
| Illness*Hispanic |  |  |  |  | 1.08 | (1.16) |  |  |
| Illness*Black |  |  |  |  | 1.24*** | (3.84) | 1.21*** | (3.91) |
| Illness*Asian |  |  |  |  | 1.06 | (0.52) |  |  |
| Illness*Other Race |  |  |  |  | 1.20 | (1.45) |  |  |
| Illness*Female |  |  |  |  | 0.89** | (-2.63) | 0.88** | (-2.70) |
| Illness*Household Income |  |  |  |  | 0.99 | (-0.21) |  |  |
| Illness*Parental Education |  |  |  |  | 0.95* | (-2.03) | 0.94** | (-2.64) |
| Illness*Asexual Attract. |  |  |  |  | 1.02 | (0.28) |  |  |
| Illness*Bisexual Attract. |  |  |  |  | 0.98 | (-0.20) |  |  |
| Illness*Same Sex Only Attract. |  |  |  |  | 1.15 | (0.91) |  |  |
| Illness*Severe Child Adversity |  |  |  |  | 0.98 | (-0.74) |  |  |
| Intercept | 50283*** | (32.78) | 47986*** | (32.95) | 47637*** | (32.98) | 47698*** | (32.96) |
| Random intercept variance | 4.41*** | (9.69) | 4.13*** | (9.44) | 4.10*** | (9.41) | 4.09*** | (9.39) |
| N | 41548 |  | 41548 |  | 41548 |  | 41548 |  |

\* $p < .05$ ; \*\* $p < .01$ ; \*\*\* $p < .001$

### Model 2: Illness Main Effects

Model 2 introduced an illness term to Model 1. The direction and magnitude of the sociodemographic, sexual attraction, and cumulative severe adversity effects remained very similar to the baseline model across all comparison. Additionally, results from all three comparisons indicated that illness was a significant risk factor for engaging in risky nonmonogamous sex. That is, for each standard deviation increase in illness, the odds of safer nonmonogamous sex decreased by 13% (*p* < .001), monogamous sex decreased by 17% (*p* < .001), and abstaining decreased by 26% (*p* < .001), compared to the risky nonmonogamous reference group.

### Model 3: Moderation of Social Disadvantage by Illness

Model 3 includes interaction effects of illness to all of the significant social factors— race/ethnicity, sex, SES, sexual attraction, and cumulative severe adversity. In the comparison of safer nonmonogamous and monogamous versus risky nonmonogamous sex (Tables 2 and 3, Model 3), the direction and magnitude of the effects of the sociodemographic variables and cumulative severe adversity remained substantively unchanged from Model 2, and no interactions were significant. In the comparison of abstinence versus risky nonmonogamous sex (Table 4, Model 3), all of the main effects for the sociodemographic variables, as well as cumulative severe adversity, and illness were substantively unchanged. Additionally, a number of interactions were significant. Being ill and having Black racial identity increased the odds of abstaining versus engaging in risky nonmonogamous sex by 24% (*p* < .001), compared to the White referent. Being both female and ill decreased the odds of abstaining (OR = 0.89, *p* < .01). This implies that the protective factor of being female is attenuated by illness, increasing the probability of engaging in risky nonmonogamous sex. Similarly, the protective factor of increased parental education is also attenuated by illness, decreasing the probability of abstaining (OR = 0.94, *p* < .05) compared to engaging in risky nonmonogamous sex.

### Model 4: Final Model

The fourth and final model included all sociodemographic variables, cumulative severe adversity, illness, and the three significant illness interactions from Model 3. All noted findings from Model 3 were substantively unchanged. Thus, in addition to significant positive associations of several social disadvantages and illness to risky nonmonogamous sex, three interactions were significant in the abstinence vs. risky nonmonogamous sex comparison (Table 4, Model 4). Hence, being both ill and female decreased the odds of abstaining by 12% (*p* < .01; see Figure 2, first panel). Similarly, being ill with higher parental educational attainment decreased the odds of abstaining by 6% (*p* < .01). Thus, again, the protective factor of higher parental education (OR = 1.154, *p* < .001) were attenuated by illness (OR = 0.789, *p* < .001; Figure 2, second panel). Finally, Black adolescents who were also ill had increased odds of abstaining by 22.5% (*p* < .001). As with the other two significant interactions visualized in Figure 2, this may be interpreted as an instance of illness attenuating the protective influence of an advantaged social status, white racial identity, in this case.

**Figure 2.**
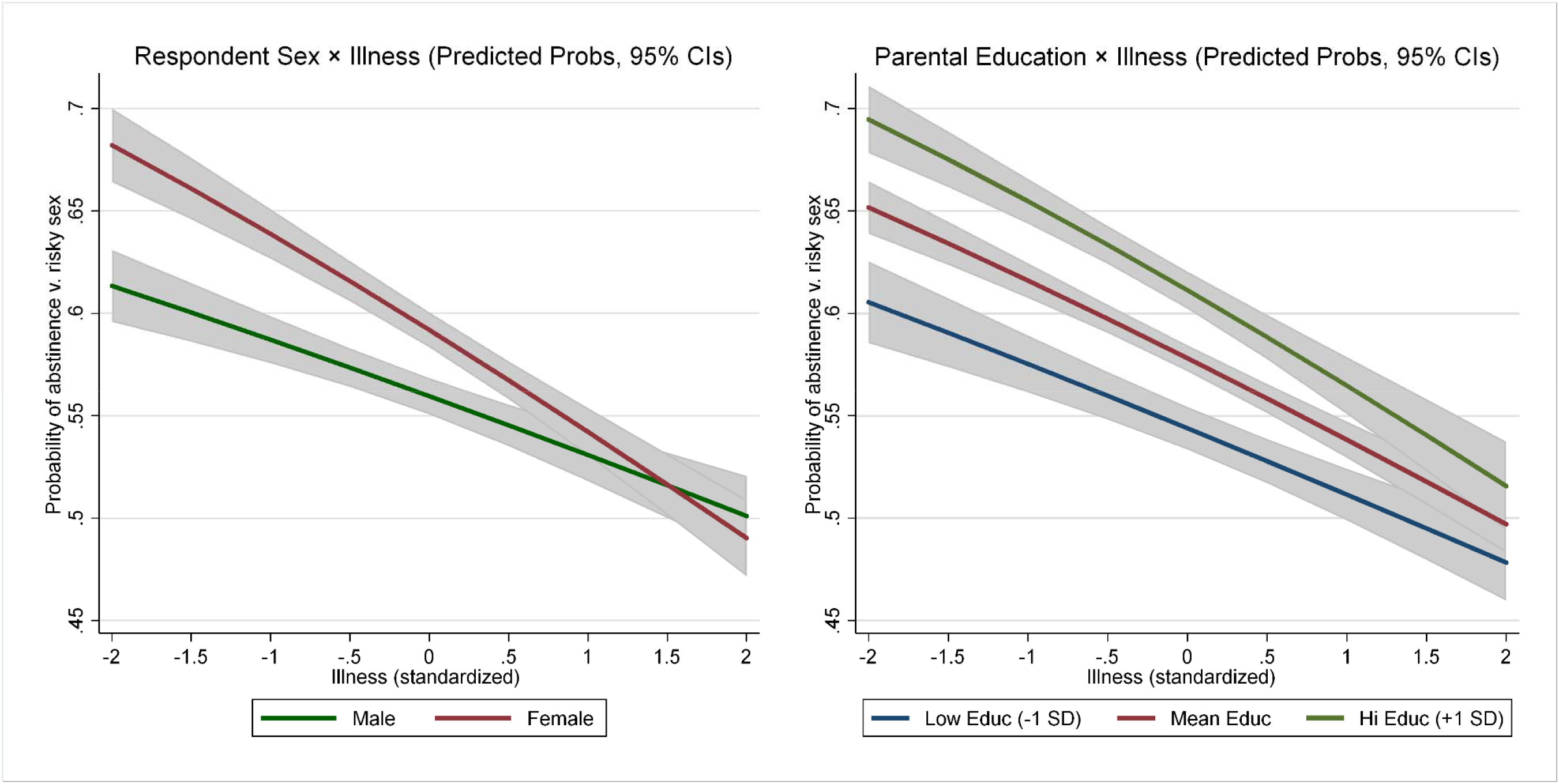
Marginal predicted probabilities of abstinence (v. risky nonmonogamous sex) visualizing respondent sex and parental education interactions with physical illness (95% CIs)

Various sensitivity analyses were conducted, and the results uniformly support the robustness of our primary analysis (see Supplemental Methods and Supplemental Table S3a-S6c for details). Specifically, sensitivity analysis comparing pooled data to wave-specific cross-sectional analyses supported robustness of the model in both mid and late adolescence (Supplemental Tables S3a-c and S4a-c); comparison of complete case analysis to MI analyses supported our use of multiple imputation (Supplemental Table S5a-c); and analyzing the five severe childhood adversity indicators separately, supported their aggregation into a cumulative index (Supplemental Table S6a-c).

## DISCUSSION

The primary aim of this study was to examine the interactive influence of social disadvantage and illness on risky sexual behavior, using the largest extant longitudinal sample of US adolescents and young adults. The analysis was guided by life history theory, which suggests that increased risk-taking may serve as an adaptive response to adversity, e.g., social disadvantage and physical illness, as individuals adapt to increased mortality risk by maximizing fitness via earlier, and riskier, sexual behavior, at the expense of longer-term parental investment strategies (Hartman, Li, Nettle, & Belsky, 2017; Hurst & Kavanagh, 2017; Simpson et al., 2012). The current results generally support these hypotheses, finding that adolescents who report higher levels of physical illness and social disadvantage are generally more likely to engage in risky nonmonogamous sex compared to safer nonmonogamous sex, monogamous sex, or abstinence. Further, results show that physical illness attenuates the protective effects of higher childhood SES, female sex, and white racial identity. Given that illness is a strong, true signal of increased mortality risk, it is, according to the logic of life history theory, unsurprising that we observe illness pervasively eroding the protective effects of social advantages.

Results are consistent with previous research demonstrating that various social disadvantages and stressors are positively associated with sexual risk-taking behavior (Hartman et al., 2017; James et al., 2012; Ryan et al., 2015; Simpson et al., 2012), supporting the hypothesis that high levels of environmental uncertainty and/or harshness promote fast life history strategies, including increased sexual risk behavior (Del Giudice et al., 2015; Szepsenwol et al., 2017). In accordance with previous research, social characteristics found to buffer against sexual risk-taking included high parental education (Del Giudice et al., 2015; Ritterman Weintraub, Fernald, Adler, Bertozzi, & Syme, 2015; Szepsenwol et al., 2017), privileged racial/ethnic identity, female sex (Clark, 1989; Grello, Welsh, & Harper, 2006). While the parental education and racial/ethnic findings likely reflect calibration of life history strategies by cumulative stress stemming from discrimination (Clark, Anderson, Clark, & Williams, 1999), stigma (Link & Phelan, 2001), and financial disadvantage (Geronimus, 1992), the female sex finding is likely due to both social control constraining young women’s sexual behavior, as well as evolutionary influences related to sex differences in parental investment (James et al., 2012; Trivers, 1972).

The second major finding in this study is the association of physical illness to risky sexual behavior, as measured by risky nonmonogamous sex, in adolescence and young adulthood. Previous research has examined the differences in sexual behaviors of adolescents with chronic conditions and disability as compared to their healthier and non-disabled peers. Much of the research conducted on this topic has found that adolescents with physical ailments engage in as much or more sex than their healthy counterparts (Cheng & Udry, 2002; Surís, Michaud, Akre, & Sawyer, 2008; Suris & Parera, 2005). LHT suggests that this is because the honest signal of physical illness on mortality risk should be a potent trigger of biobehavioral adaptation toward fast life history strategies, particularly in adolescence, as the cusp of sexuality and fertility (Anderson, 2017; Chisholm et al., 1993). That is not to say, however, that there is a general association of health conditions to increased sexual activity. For instance, Kahn and Halpern recently found that among individuals in their 30’s, those with physical disabilities were less likely to report having sex compared to those without disabilities (Kahn & Halpern, 2018). Future research will be needed to integrate these findings, and contrasting the current study with Kahn and Halpern (2018) indicates several potentially high impact topics for future research including exploring: (a) the differential impacts of disability and illness; (b) possible heterogeneity in our risky nonmonogamous sex findings across developmental periods (e.g., adolescence versus mature adulthood); and (c) disaggregating the sexually active population into more granular categories.

Notably, this study found a coherent pattern of attenuation of the protective effects of social advantages by physical illness. Thus, across race/ethnicity, sex, and socioeconomic statuses, the protective effects of social advantage are progressively eroded as physical illness increases. These findings generally supported expectations, although the sex × illness interaction, in particular, certainly warrants further research, due to opposing theoretical expectations within evolutionary psychology regarding the expected impact of adversity on male sexual behavior (James et al., 2012). Specifically, it has been argued that given baseline sex differences in sexual behavior (Trivers, 1972), illness may be expected to reduce short-term mating opportunities, and consequently, sexual risk behavior in males. Future research will be needed to adjudicate this question more definitively, and it may well be that the effect of illness of sexual risk behavior depends on the developmental stage. Despite the need for additional research to replicate and extend these findings, the high levels of statistical significance (generally, *p* < .001) and robustness to various sensitivity analyses (see Supplemental Material) indicate that the influence of physical illness promoting sexual risk behavior is pervasive, interactive, and powerful, at least during adolescence and the transition to adulthood.

While we generally frame this analysis in the life history perspective, the findings support a complex, nuanced interpretation, including the influence of cultural and structural social factors. For example, our finding that first-generation immigrants were less likely to engage in risky nonmonogamous sex than their native peers contradicts the naïve LHT prediction that individuals facing increased social adversity (i.e., first-generation immigrants) will be more likely to engage in risky sexual behavior. Clearly, cultural factors are at play in this instance. Structural inequalities are likely driving other results, particularly the observed racial/ethnic differences. That is, the structurally maintained experience of discrimination, and attendant stressors, likely functions as a significant calibrating adversity for disadvantaged racial/ethnic groups. This suggests that a fruitful synthesis may be achieved by integrating life history perspectives with established structural inequality paradigms regarding racial/ethnic discrimination (Williams & Mohammed, 2009; Hicken, Lee, Morenoff, House, & Williams, 2014; Clark et al. 1999), minority stress theory (Meyer, 2003), stigma (Link & Phelan, 2001), and the weathering hypothesis (Geronimus, 1992).

Notwithstanding its merits, the current study is characterized by various limitations. Due to the nature of retrospective self-report questionnaires, the study is vulnerable to limitations such as recall bias, which may influence results. This is a universal limitation of observational research; however, relatively short windows for retrospection, a very large sample size, and repeated assessments per individual ensure that the current study does not suffer disproportionately from this limitation. Another noteworthy limitation stems from the trade-off between parsimony and comprehensiveness in our specification of the sexual behavior outcome. That is, to keep our study tightly focused on sexual risk, we limited our classification of sexual behavior to four categories. Future research should consider a broader range of meaningful subdivisions, more fully articulating the diverse identities, experiences, and behaviors within nonmonogamous populations, including parsing subgroups based on infidelity, number of partners, and casual hookups versus stable polyamorous relationships. Finally, we explicitly note that the current study, though theoretically informed, was exploratory and not pre-registered. We encourage future efforts to replicate this study to pre-register to support unbiased assessment of the robustness of the findings.

Despite these limitations, the current study advances understanding of how illness impacts risk behavior, and notably supports and extends life history theorizing regarding sexual behavior trade-offs in adolescence (Ellis & Del Giudice, 2014; Ellis et al., 2012). These findings suggest the need for further research exploring the relationship between physical illness and risk behaviors across the life course. Additionally, future work in this area should consider mental health and specific childhood disease states, as well as further explore variation in LH-sexuality linkages across sexual orientation. Developing more accurate and nuanced models of adolescent sexual risk can identify promising areas for intervention by parents, healthcare providers, and policymakers to support health and well-being in adolescence and the transition to adulthood.

### Ethical approval

All procedures performed in studies involving human participants were in accordance with the ethical standards of the institutional and/or national research committee (University of Utah, Institutional Review Board; IRB_00107767) and with the 1964 Helsinki declaration and its later amendments or comparable ethical standards.

## Acknowledgements

We thank the Consortium of Families and Health Research (C-FAHR) at the University of Utah for providing the forum in which this paper was developed. We are grateful to Cynthia A. Berg for editorial input, Claudia Geist for methodological advice, and Bruce Ellis for intellectual inspiration. This research uses data from Add Health, a program project directed by Kathleen Mullan Harris and designed by J. Richard Udry, Peter S. Bearman, and Kathleen Mullan Harris at the University of North Carolina at Chapel Hill, and funded by grant P01-HD31921 from the Eunice Kennedy Shriver National Institute of Child Health and Human Development, with cooperative funding from 23 other federal agencies and foundations. Special acknowledgment is due Ronald R. Rindfuss and Barbara Entwisle for assistance in the original design. Information on how to obtain the Add Health data files is available on the Add Health website (http://www.cpc.unc.edu/addhealth). No direct support was received from grant P01-HD31921 for this analysis.

