## Supplementary Material for "Illness, social disadvantage, and sexual risk behavior in adolescence and the transition to adulthood"

**Supplemental Methods**

*Severe childhood adversity*

Measures of childhood adversity comprised five dichotomous indicators: *sexual abuse, physical abuse, neglect, drugs/alcohol accessible at home, and parental incarceration,* occurring in adolescence. These five severe childhood adversities were operationalized as follows (item phrasing condensed for brevity). *Sexual abuse* was indicated by endorsement of any of the following four items: the respondent sexually interacted with a parent or other adult caregiver by grade 6; the respondent sexually interacted with a parent or other adult caregiver as a minor; or either of two items asking if the respondent had been sexually coerced as a minor, either physically, or non-physically (e.g., drugged). *Physical abuse* was indicated by endorsement of either of two items: the respondent was slapped, hit, or kicked by parents or other adult care-givers by grade 6; or as a minor. *Neglect* was indicated by endorsement of any of the following items: the respondent’s parent/s or other adult caregiver/s had not taken care of basic needs, such as keeping the respondent clean or providing food or clothing by grade 6; respondent frequently felt unloved or unwanted by parent/s or caregiver/s as a minor. *Drugs/alcohol accessible at home* was indicated by endorsement of any of the following items: the respondent had easy access to illegal drugs in the home; respondent reported frequent heavy drinking in the home (e.g., open containers, intoxicated people). *Parental incarceration* was indicated by endorsement of any of the following items: the respondent’s biological father, biological mother, “father figure”, or “mother figure” went to jail/prison, when respondent was a minor. A cumulative adversity count was used ranging from 0 to 5.

*Sociodemographic characteristics*

All presented models included a set of sociodemographic covariates characterizing the adolescent’s social background. These included household SES measures for parental education and household income assessed at Wave 1. We used the parental questionnaire to ascertain the educational attainment of the sampled residential parent (typically mother) and current partner (typically father). Parental educational attainments were averaged across parents when more than one was reported. Household income was ascertained from the parental questionnaire and included all sources of income from the previous year (measured in thousands of dollars) and is logged (to reduce pronounced positive skew). Both parental education and income were standardized in the analysis. Self-reported race/ethnicity (coded Black, White (ref), Asian, Hispanic, and Other), nativity (i.e., first-generation immigrant status), and age were also included as covariates in all presented models.

*Multiple imputation*

There was a small-to-moderate amount of missing data at both waves. The amount of missing data was modest for the time invariant variables with only household income (26.87%), sexual abuse (25.20%), physical abuse (15.07%), parental incarceration (24.42%), and neglect (28.03%) exceeding >5% missing. Among the time variant variables, sexual risk status (i.e., casual/relationship-only/abstinent) and the physical illness construct both had little missingness at Wave 1 (>2%) and a moderate amount of missingness (~29%) in Wave 2 (see Supplemental Table S2 for full missingness information). To avoid the potential of biased estimation due to scenarios of missing completely at random and missingness at random, we addressed missing data using multiple imputation (MI).*^13^* As the missing data pattern was non-monotone, we used *multiple imputation with chained equations* (*MICE*; i.e., fully conditional specification or sequential generalized regression) as implemented in Stata 15.1. This is a well-validated MI method, which has been implemented in Stata for several versions. The primary advantage of MICE over other MI methods is its ability to maintain the distributional characteristics of the imputed variables. Thus, in our MICE data generation model we specified sex, the five severe adversities, the sexual behavior outcomes, and sexual attraction items using the logit link function, and treated the other variables using the standard (continuous) linear link function. Care was taken to capitalize on the longitudinal information in the data in the imputation, by organizing the time variant variables wide for the imputation and then reshaping the data long for the analysis, using Stata’s MI data management facilities. We used a conservative 30 imputations^16,17^ in all MI analyses, which were conducted using the “mi estimate” prefix command in Stata. The “mi estimate” runs estimation commands (mixed effects multinomial logistic regression, in this case) on the imputed MI data, and adjusts coefficients and standard errors for the variability between imputations according to the combination rules by Rubin^18^.

*Sensitivity analysis*

To test the robustness of our substantive results we conducted several sensitivity analyses. These results support our use of multiple imputation, pooled data, and combining childhood adversity indicators into a cumulative measure. Sensitivity analyses include the following: (A) the primary model sequence stratified by wave. Supplemental Tables S3a-c and S4a-c show the cross-sectional results stratified by wave. While some associations change slightly in the relationship-only versus casual sex comparison in Wave I, these differences are modest and generally not present in Wave II, when compared to the pooled results. The results of the abstinence versus nonmonogamous sex comparison are substantively identical across the wave-specific analyses and the pooled results. (B) The primary model sequence using complete case analysis rather than multiple imputation (Supplemental Tables S5a-c). Comparison of the results reported using multiple imputation (Tables 2-4) to those using complete case indicates that, while the results are substantively very similar, the magnitude of the statistical associations is generally increased by the use of multiple imputation. (C) A model substituting the five individual severe childhood adversities (i.e., dichotomous indicators of sexual abuse, physical abuse, neglect, drugs/alcohol in the home, parental incarceration) for the cumulative childhood adversity index (Supplemental Tables S6a-c). The results indicate that while each adversity is independently significantly positively associated with nonmonogamous sex, the cumulative specification has stronger predictive power than any of the five indicators individually, thus supporting its use in the main text.

**Table S2.** Missingness by analysis variable. Time variant measures indicated by wave number

| **Variable** | **Missing (obs)** | **Total** | **Missing (%)** |
| --- | --- | --- | --- |
| Wave 1: Sexual risk behavior | 336 | 20,774 | 1.62 |
| Wave 2: Sexual risk behavior | 6,167 | 20,774 | 29.69 |
| Wave 1: Self-rated illness | 55 | 20,774 | 0.26 |
| Wave 2: Self-rated illness | 6,040 | 20,774 | 29.07 |
| Wave 1: Sexual orientation | 299 | 20,774 | 1.44 |
| Wave 2: Sexual orientation | 6,135 | 20,774 | 29.53 |
| Wave 1: Age | 46 | 20,774 | 0.22 |
| Wave 2: Age | 32 | 20,774 | 0.15 |
| Immigration status | 0 | 20,774 | 0 |
| Race/ethnicity | 0 | 20,774 | 0 |
| Sex | 31 | 20,774 | 0.15 |
| Household Income | 5,582 | 20,774 | 26.87 |
| Parental Education | 759 | 20,774 | 3.65 |
| Sexual abuse (adversity index item) | 5,236 | 20,774 | 25.2 |
| Physical abuse (adv. index item) | 3,130 | 20,774 | 15.07 |
| Neglect (adv. index item) | 5,823 | 20,774 | 28.03 |
| Drugs/alcohol in home (adv. index item) | 32 | 20,774 | 0.15 |
| Parental incarceration (adv. index item) | 5,073 | 20,774 | 24.42 |

**Table S3a.** Main analysis stratified by Wave, Wave 1; Protected nonmonogamous sex v. risky nonmonogamous sex comparison (MI, m=30)

|  | **Model 1** | | **Model 2** | | **Model 3** | | **Model 4** | |
| --- | --- | --- | --- | --- | --- | --- | --- | --- |
| **Main effects** | **OR** | **t** | **OR** | **t** | **OR** | **t** | **OR** | **t** |
| Immigrant | 1.25 | (1.51) | 1.22 | (1.32) | 1.22 | (1.33) | 1.22 | (1.33) |
| Hispanic | 0.79* | (-2.57) | 0.79* | (-2.52) | 0.78** | (-2.58) | 0.79* | (-2.51) |
| Black | 1.08 | (1.08) | 1.06 | (0.85) | 1.05 | (0.62) | 1.05 | (0.67) |
| Asian | 0.53*** | (-3.94) | 0.55*** | (-3.66) | 0.54*** | (-3.64) | 0.56*** | (-3.62) |
| Other Race | 0.80 | (-1.33) | 0.81 | (-1.26) | 0.79 | (-1.37) | 0.81 | (-1.25) |
| Female | 0.77*** | (-4.50) | 0.79*** | (-3.95) | 0.79*** | (-3.80) | 0.79*** | (-3.82) |
| Household Income | 1.07 | (1.83) | 1.07 | (1.74) | 1.07 | (1.82) | 1.07 | (1.70) |
| Parental Education | 1.03 | (0.78) | 1.02 | (0.44) | 1.02 | (0.56) | 1.02 | (0.66) |
| Age | 0.94*** | (-3.35) | 0.93*** | (-3.39) | 0.93*** | (-3.37) | 0.93*** | (-3.40) |
| Asexual Attraction | 0.85 | (-1.19) | 0.86 | (-1.15) | 0.84 | (-1.25) | 0.86 | (-1.16) |
| Bisexual Attraction | 0.89 | (-1.04) | 0.90 | (-0.94) | 0.92 | (-0.73) | 0.90 | (-0.93) |
| Same Sex Only Attract. | 0.83 | (-0.76) | 0.86 | (-0.65) | 0.83 | (-0.73) | 0.85 | (-0.68) |
| Severe Child Adversity | 0.90** | (-3.27) | 0.91** | (-2.96) | 0.91** | (-2.81) | 0.91** | (-2.94) |
| Illness |  |  | 0.88*** | (-4.42) | 0.85** | (-2.83) | 0.85*** | (-3.73) |
| **Interaction effects** |  |  |  |  |  |  |  |  |
| Illness*Immigrant |  |  |  |  | 0.94 | (-0.44) |  |  |
| Illness*Hispanic |  |  |  |  | 1.00 | (-0.02) |  |  |
| Illness*Black |  |  |  |  | 1.00 | (0.06) | 1.02 | (0.28) |
| Illness*Asian |  |  |  |  | 1.07 | (0.44) |  |  |
| Illness*Other Race |  |  |  |  | 1.08 | (0.48) |  |  |
| Illness*Female |  |  |  |  | 1.07 | (1.22) | 1.07 | (1.23) |
| Illness*Household Income |  |  |  |  | 0.97 | (-0.94) |  |  |
| Illness*Parental Education |  |  |  |  | 0.99 | (-0.30) | 0.98 | (-0.82) |
| Illness*Asexual Attract. |  |  |  |  | 1.10 | (0.79) |  |  |
| Illness*Bisexual Attract. |  |  |  |  | 0.92 | (-0.72) |  |  |
| Illness*Same Sex Only Attract. |  |  |  |  | 1.09 | (0.42) |  |  |
| Illness*Severe Child Adversity |  |  |  |  | 1.00 | (-0.17) |  |  |
| Intercept | 4.57*** | (4.37) | 4.60*** | (4.38) | 4.56*** | (4.35) | 4.60*** | (4.38) |
| N | 20774 | | 20774 | | 20774 | | 20774 | |
| * p<0.05; ** p<0.01; *** p<0.001 | |  |  |  |  |  |  |  |

**Table S3b.** Main analysis stratified by Wave, Wave 1; Monogamous sex v. risky nonmonogamous sex comparison (MI, m=30)

|  | **Model 1** | | **Model 2** | | **Model 3** | | **Model 4** | |
| --- | --- | --- | --- | --- | --- | --- | --- | --- |
| **Main effects** | **OR** | **t** | **OR** | **t** | **OR** | **t** | **OR** | **t** |
| Immigrant | 1.54*** | (3.42) | 1.48** | (3.10) | 1.49** | (3.14) | 1.48** | (3.09) |
| Hispanic | 0.98 | (-0.30) | 0.98 | (-0.21) | 0.96 | (-0.50) | 0.98 | (-0.20) |
| Black | 0.83** | (-2.72) | 0.81** | (-3.06) | 0.79*** | (-3.30) | 0.80** | (-3.21) |
| Asian | 0.83 | (-1.43) | 0.88 | (-0.96) | 0.86 | (-1.08) | 0.89 | (-0.85) |
| Other Race | 0.73 | (-1.96) | 0.74 | (-1.83) | 0.72 | (-1.95) | 0.74 | (-1.83) |
| Female | 1.27*** | (4.34) | 1.33*** | (5.12) | 1.35*** | (5.34) | 1.35*** | (5.31) |
| Household Income | 1.00 | (0.03) | 1.00 | (-0.13) | 1.00 | (-0.04) | 0.99 | (-0.15) |
| Parental Education | 1.04 | (1.31) | 1.03 | (0.78) | 1.03 | (0.88) | 1.03 | (1.00) |
| Age | 0.92*** | (-4.21) | 0.92*** | (-4.28) | 0.92*** | (-4.23) | 0.92*** | (-4.27) |
| Asexual Attraction | 1.44** | (3.16) | 1.45** | (3.23) | 1.43** | (3.09) | 1.45** | (3.22) |
| Bisexual Attraction | 0.81 | (-1.94) | 0.82 | (-1.80) | 0.84 | (-1.49) | 0.83 | (-1.73) |
| Same Sex Only Attract. | 0.71 | (-1.52) | 0.74 | (-1.34) | 0.75 | (-1.22) | 0.74 | (-1.32) |
| Severe Child Adversity | 0.85*** | (-5.62) | 0.87*** | (-5.10) | 0.87*** | (-4.83) | 0.87*** | (-5.09) |
| Illness |  |  | 0.83*** | (-7.02) | 0.82*** | (-3.53) | 0.84*** | (-4.18) |
| **Interaction effects** |  |  |  |  |  |  |  |  |
| Illness*Immigrant |  |  |  |  | 0.99 | (-0.08) |  |  |
| Illness*Hispanic |  |  |  |  | 1.10 | (1.21) |  |  |
| Illness*Black |  |  |  |  | 1.08 | (1.16) | 1.06 | (0.98) |
| Illness*Asian |  |  |  |  | 1.09 | (0.66) |  |  |
| Illness*Other Race |  |  |  |  | 1.11 | (0.67) |  |  |
| Illness*Female |  |  |  |  | 0.93 | (-1.45) | 0.92 | (-1.49) |
| Illness*Household Income |  |  |  |  | 0.98 | (-0.64) |  |  |
| Illness*Parental Education |  |  |  |  | 0.97 | (-0.89) | 0.96 | (-1.69) |
| Illness*Asexual Attract. |  |  |  |  | 1.08 | (0.77) |  |  |
| Illness*Bisexual Attract. |  |  |  |  | 0.87 | (-1.29) |  |  |
| Illness*Same Sex Only Attract. |  |  |  |  | 0.99 | (-0.03) |  |  |
| Illness*Severe Child Adversity |  |  |  |  | 0.99 | (-0.47) |  |  |
| Intercept | 6.49*** | (5.78) | 6.56*** | (5.80) | 6.49*** | (5.76) | 6.54*** | (5.79) |
| N | 20774 | | 20774 | | 20774 | | 20774 | |
| * p<0.05; ** p<0.01; *** p<0.001 | |  |  |  |  |  |  |  |

**Table S3c.** Main analysis stratified by Wave, Wave 1; Abstinent v. risky nonmonogamous sex comparison (MI, m=30)

|  | **Model 1** | | **Model 2** | | | **Model 3** | | | **Model 4** | | |
| --- | --- | --- | --- | --- | --- | --- | --- | --- | --- | --- | --- |
| **Main effects** | **OR** | **t** | | **OR** | **t** | | **OR** | **t** | | **OR** | **t** |
| Immigrant | 2.786*** | (8.95) | | 2.635*** | (8.43) | | 2.627*** | (8.34) | | 2.625*** | (8.39) |
| Hispanic | 0.928 | (-1.00) | | 0.940 | (-0.83) | | 0.942 | (-0.80) | | 0.942 | (-0.80) |
| Black | 0.408*** | (-14.16) | | 0.395*** | (-14.60) | | 0.395*** | (-14.53) | | 0.396*** | (-14.51) |
| Asian | 1.141 | (1.14) | | 1.245 | (1.88) | | 1.231 | (1.71) | | 1.273* | (2.07) |
| Other Race | 0.681** | (-2.75) | | 0.699* | (-2.55) | | 0.686** | (-2.60) | | 0.700* | (-2.53) |
| Female | 1.219*** | (3.98) | | 1.298*** | (5.19) | | 1.312*** | (5.31) | | 1.312*** | (5.32) |
| Household Income | 1.121*** | (3.49) | | 1.110** | (3.19) | | 1.109** | (3.15) | | 1.109** | (3.14) |
| Parental Education | 1.219*** | (6.63) | | 1.190*** | (5.79) | | 1.196*** | (5.89) | | 1.196*** | (5.90) |
| Age | 0.546*** | (-35.78) | | 0.545*** | (-35.76) | | 0.545*** | (-35.72) | | 0.545*** | (-35.74) |
| Asexual Attraction | 3.034*** | (10.84) | | 3.066*** | (10.91) | | 3.017*** | (10.69) | | 3.053*** | (10.88) |
| Bisexual Attraction | 0.722** | (-3.22) | | 0.737** | (-3.00) | | 0.760** | (-2.63) | | 0.747** | (-2.87) |
| Same Sex Only Attract. | 0.783 | (-1.22) | | 0.823 | (-0.96) | | 0.844 | (-0.79) | | 0.829 | (-0.92) |
| Severe Child Adversity | 0.637*** | (-16.91) | | 0.650*** | (-16.15) | | 0.652*** | (-15.66) | | 0.650*** | (-16.12) |
| Illness |  |  | | 0.762*** | (-11.15) | | 0.759*** | (-6.04) | | 0.763*** | (-7.18) |
| **Interaction effects** |  |  | |  |  | |  |  | |  |  |
| Illness*Immigrant |  |  | |  |  | | 0.96 | (-0.34) | |  |  |
| Illness*Hispanic |  |  | |  |  | | 1.13 | (1.67) | |  |  |
| Illness*Black |  |  | |  |  | | 1.25*** | (3.61) | | 1.20*** | (3.43) |
| Illness*Asian |  |  | |  |  | | 1.16 | (1.30) | |  |  |
| Illness*Other Race |  |  | |  |  | | 1.11 | (0.76) | |  |  |
| Illness*Female |  |  | |  |  | | 0.89* | (-2.34) | | 0.89* | (-2.46) |
| Illness*Household Income |  |  | |  |  | | 0.99 | (-0.45) | |  |  |
| Illness*Parental Education |  |  | |  |  | | 0.96 | (-1.56) | | 0.94* | (-2.38) |
| Illness*Asexual Attract. |  |  | |  |  | | 1.06 | (0.66) | |  |  |
| Illness*Bisexual Attract. |  |  | |  |  | | 0.99 | (-0.15) | |  |  |
| Illness*Same Sex Only Attract. |  |  | |  |  | | 0.92 | (-0.44) | |  |  |
| Illness*Severe Child Adversity |  |  | |  |  | | 0.99 | (-0.51) | |  |  |
| Intercept | 154031*** | (40.91) | | 155442*** | (40.79) | | 155831*** | (40.74) | | 155290*** | (40.77) |
| N | 20774 | | 20774 | | | 20774 | | | 20774 | | |
| * p<0.05; ** p<0.01; *** p<0.001 | |  |  | |  |  | |  |  | |  |

**Table S4a.** Main analysis stratified by Wave, Wave 2; Protected nonmonogamous sex v. risky nonmonogamous sex comparison (MI, m=30)

|  | **Model 1** | | **Model 2** | | **Model 3** | | **Model 4** | |
| --- | --- | --- | --- | --- | --- | --- | --- | --- |
| **Main effects** | **OR** | **t** | **OR** | **t** | **OR** | **t** | **OR** | **t** |
| Immigrant | 0.62* | (-2.21) | 0.60* | (-2.33) | 0.61* | (-2.22) | 0.60* | (-2.35) |
| Hispanic | 0.80 | (-1.66) | 0.80 | (-1.63) | 0.81 | (-1.50) | 0.80 | (-1.63) |
| Black | 0.80* | (-2.20) | 0.79* | (-2.34) | 0.78* | (-2.41) | 0.78* | (-2.43) |
| Asian | 0.50** | (-2.83) | 0.52** | (-2.72) | 0.54* | (-2.35) | 0.52** | (-2.67) |
| Other Race | 0.96 | (-0.16) | 0.97 | (-0.13) | 0.92 | (-0.35) | 0.97 | (-0.12) |
| Female | 0.93 | (-0.77) | 0.96 | (-0.47) | 0.97 | (-0.37) | 0.97 | (-0.35) |
| Household Income | 1.10 | (1.91) | 1.10 | (1.84) | 1.09 | (1.79) | 1.10 | (1.83) |
| Parental Education | 1.14** | (2.63) | 1.13* | (2.47) | 1.14* | (2.57) | 1.14* | (2.53) |
| Age | 1.06 | (1.48) | 1.06 | (1.49) | 1.06 | (1.49) | 1.06 | (1.49) |
| Asexual Attraction | 0.87 | (-0.93) | 0.88 | (-0.91) | 0.88 | (-0.86) | 0.87 | (-0.92) |
| Bisexual Attraction | 0.82 | (-1.12) | 0.83 | (-1.06) | 0.80 | (-1.22) | 0.83 | (-1.05) |
| Same Sex Only Attract. | 0.54 | (-1.47) | 0.54 | (-1.46) | 0.53 | (-1.49) | 0.55 | (-1.45) |
| Severe Child Adversity | 0.92 | (-1.70) | 0.93 | (-1.57) | 0.93 | (-1.45) | 0.93 | (-1.53) |
| Illness |  |  | 0.91** | (-2.63) | 0.90 | (-1.38) | 0.88* | (-2.03) |
| **Interaction effects** |  |  |  |  |  |  |  |  |
| Illness*Immigrant |  |  |  |  | 0.87 | (-0.67) |  |  |
| Illness*Hispanic |  |  |  |  | 0.93 | (-0.60) |  |  |
| Illness*Black |  |  |  |  | 1.10 | (0.99) | 1.11 | (1.25) |
| Illness*Asian |  |  |  |  | 0.83 | (-0.85) |  |  |
| Illness*Other Race |  |  |  |  | 1.27 | (1.16) |  |  |
| Illness*Female |  |  |  |  | 0.94 | (-0.66) | 0.95 | (-0.58) |
| Illness*Household Income |  |  |  |  | 1.01 | (0.32) |  |  |
| Illness*Parental Education |  |  |  |  | 0.95 | (-1.13) | 0.96 | (-0.88) |
| Illness*Asexual Attract. |  |  |  |  | 0.96 | (-0.32) |  |  |
| Illness*Bisexual Attract. |  |  |  |  | 1.17 | (0.96) |  |  |
| Illness*Same Sex Only Attract. |  |  |  |  | 1.65 | (1.40) |  |  |
| Illness*Severe Child Adversity |  |  |  |  | 0.99 | (-0.17) |  |  |
| Intercept | 0.58 | (-0.85) | 0.58 | (-0.87) | 0.58 | (-0.86) | 0.58 | (-0.87) |
| N | 20774 | | 20774 | | 20774 | | 20774 | |
| * p<0.05; ** p<0.01; *** p<0.001 | |  |  |  |  |  |  |  |

**Table S4b.** Main analysis stratified by Wave, Wave 2; Monogamous sex v. risky nonmonogamous sex comparison (MI, m=30)

|  | **Model 1** | | **Model 2** | | **Model 3** | | **Model 4** | |
| --- | --- | --- | --- | --- | --- | --- | --- | --- |
| **Main effects** | **OR** | **t** | **OR** | **t** | **OR** | **t** | **OR** | **t** |
| Immigrant | 0.910 | (-0.56) | 0.878 | (-0.78) | 0.899 | (-0.63) | 0.876 | (-0.80) |
| Hispanic | 0.911 | (-0.91) | 0.917 | (-0.85) | 0.916 | (-0.86) | 0.917 | (-0.85) |
| Black | 0.587*** | (-6.25) | 0.576*** | (-6.48) | 0.568*** | (-6.64) | 0.570*** | (-6.61) |
| Asian | 0.911 | (-0.54) | 0.945 | (-0.33) | 0.975 | (-0.13) | 0.954 | (-0.27) |
| Other Race | 0.765 | (-1.28) | 0.773 | (-1.23) | 0.739 | (-1.46) | 0.776 | (-1.22) |
| Female | 1.680*** | (7.33) | 1.743*** | (7.72) | 1.759*** | (7.64) | 1.761*** | (7.64) |
| Household Income | 1.031 | (0.73) | 1.026 | (0.61) | 1.025 | (0.59) | 1.025 | (0.59) |
| Parental Education | 1.140** | (3.10) | 1.127** | (2.83) | 1.134** | (2.96) | 1.132** | (2.90) |
| Age | 1.084** | (2.96) | 1.084** | (2.97) | 1.084** | (2.97) | 1.084** | (2.97) |
| Asexual Attraction | 2.115*** | (7.35) | 2.125*** | (7.39) | 2.139*** | (7.36) | 2.124*** | (7.39) |
| Bisexual Attraction | 0.539*** | (-3.95) | 0.548*** | (-3.84) | 0.549*** | (-3.73) | 0.548*** | (-3.84) |
| Same Sex Only Attract. | 1.104 | (0.36) | 1.106 | (0.36) | 1.114 | (0.38) | 1.111 | (0.38) |
| Severe Child Adversity | 0.870*** | (-4.01) | 0.877*** | (-3.76) | 0.883*** | (-3.47) | 0.878*** | (-3.71) |
| Illness |  |  | 0.870*** | (-4.35) | 0.868* | (-2.39) | 0.840*** | (-3.57) |
| **Interaction effects** |  |  |  |  |  |  |  |  |
| Illness*Immigrant |  |  |  |  | 0.81 | (-1.56) |  |  |
| Illness*Hispanic |  |  |  |  | 0.98 | (-0.17) |  |  |
| Illness*Black |  |  |  |  | 1.08 | (0.96) | 1.09 | (1.22) |
| Illness*Asian |  |  |  |  | 0.98 | (-0.11) |  |  |
| Illness*Other Race |  |  |  |  | 1.22 | (1.11) |  |  |
| Illness*Female |  |  |  |  | 0.98 | (-0.32) | 0.98 | (-0.31) |
| Illness*Household Income |  |  |  |  | 1.00 | (-0.02) |  |  |
| Illness*Parental Education |  |  |  |  | 0.97 | (-0.95) | 0.98 | (-0.76) |
| Illness*Asexual Attract. |  |  |  |  | 0.96 | (-0.45) |  |  |
| Illness*Bisexual Attract. |  |  |  |  | 0.99 | (-0.09) |  |  |
| Illness*Same Sex Only Attract. |  |  |  |  | 1.24 | (0.83) |  |  |
| Illness*Severe Child Adversity |  |  |  |  | 0.98 | (-0.85) |  |  |
| Intercept | 0.97 | (-0.08) | 0.96 | (-0.10) | 0.95 | (-0.11) | 0.95 | (-0.11) |
| N | 20774 | | 20774 | | 20774 | | 20774 | |
| * p<0.05; ** p<0.01; *** p<0.001 | |  |  |  |  |  |  |  |

**Table S4c.** Main analysis stratified by Wave, Wave 2; Abstinent v. risky nonmonogamous sex comparison (MI, m=30)

|  | **Model 1** | | **Model 2** | | | **Model 3** | | | **Model 4** | | |
| --- | --- | --- | --- | --- | --- | --- | --- | --- | --- | --- | --- |
| **Main effects** | **OR** | **t** | | **OR** | **t** | | **OR** | **t** | | **OR** | **t** |
| Immigrant | 1.67** | (3.24) | | 1.56** | (2.79) | | 1.59** | (2.83) | | 1.55** | (2.78) |
| Hispanic | 0.88 | (-1.24) | | 0.89 | (-1.13) | | 0.89 | (-1.05) | | 0.89 | (-1.12) |
| Black | 0.36*** | (-11.83) | | 0.35*** | (-12.12) | | 0.35*** | (-12.05) | | 0.35*** | (-12.09) |
| Asian | 1.50* | (2.45) | | 1.61** | (2.88) | | 1.68** | (2.89) | | 1.64** | (2.99) |
| Other Race | 0.69 | (-1.91) | | 0.70 | (-1.81) | | 0.67* | (-2.02) | | 0.70 | (-1.78) |
| Female | 1.45*** | (5.66) | | 1.55*** | (6.52) | | 1.55*** | (6.41) | | 1.55*** | (6.43) |
| Household Income | 1.15*** | (3.43) | | 1.14** | (3.14) | | 1.14** | (3.06) | | 1.14** | (3.12) |
| Parental Education | 1.28*** | (5.85) | | 1.25*** | (5.24) | | 1.25*** | (5.22) | | 1.25*** | (5.19) |
| Age | 0.65*** | (-17.62) | | 0.65*** | (-17.75) | | 0.65*** | (-17.71) | | 0.65*** | (-17.68) |
| Asexual Attraction | 2.79*** | (10.12) | | 2.81*** | (10.16) | | 2.82*** | (10.14) | | 2.80*** | (10.16) |
| Bisexual Attraction | 0.55*** | (-4.00) | | 0.57*** | (-3.78) | | 0.57*** | (-3.67) | | 0.58*** | (-3.76) |
| Same Sex Only Attract. | 1.08 | (0.28) | | 1.08 | (0.29) | | 1.10 | (0.35) | | 1.09 | (0.33) |
| Severe Child Adversity | 0.63*** | (-13.45) | | 0.64*** | (-12.88) | | 0.65*** | (-12.33) | | 0.65*** | (-12.76) |
| Illness |  |  | | 0.76*** | (-8.67) | | 0.76*** | (-4.38) | | 0.75*** | (-5.92) |
| **Interaction effects** |  |  | |  |  | |  |  | |  |  |
| Illness*Immigrant |  |  | |  |  | | 0.84 | (-1.36) | |  |  |
| Illness*Hispanic |  |  | |  |  | | 1.01 | (0.12) | |  |  |
| Illness*Black |  |  | |  |  | | 1.23** | (2.61) | | 1.22** | (2.86) |
| Illness*Asian |  |  | |  |  | | 0.97 | (-0.21) | |  |  |
| Illness*Other Race |  |  | |  |  | | 1.23 | (1.14) | |  |  |
| Illness*Female |  |  | |  |  | | 0.89 | (-1.78) | | 0.89 | (-1.78) |
| Illness*Household Income |  |  | |  |  | | 1.01 | (0.38) | |  |  |
| Illness*Parental Education |  |  | |  |  | | 0.94 | (-1.66) | | 0.95 | (-1.64) |
| Illness*Asexual Attract. |  |  | |  |  | | 0.98 | (-0.18) | |  |  |
| Illness*Bisexual Attract. |  |  | |  |  | | 1.07 | (0.46) | |  |  |
| Illness*Same Sex Only Attract. |  |  | |  |  | | 1.66* | (2.02) | |  |  |
| Illness*Severe Child Adversity |  |  | |  |  | | 0.99 | (-0.36) | |  |  |
| Intercept | 16061*** | (22.73) | | 15502*** | (22.79) | | 15529*** | (22.74) | | 15483*** | (22.72) |
| N | 20774 | | 20774 | | | 20774 | | | 20774 | | |
| * p<0.05; ** p<0.01; *** p<0.001 | |  |  | |  |  | |  |  | |  |

**Table S5a.** Main analysis using complete case analysis (rather than multiple imputation); Protected nonmonogamous sex v. risky nonmonogamous sex comparison

|  | **Model 1** | | **Model 2** | | **Model 3** | | **Model 4** | |
| --- | --- | --- | --- | --- | --- | --- | --- | --- |
| **Main effects** | **OR** | **t** | **OR** | **t** | **OR** | **t** | **OR** | **t** |
| Immigrant | 1.16 | (0.47) | 1.10 | (0.31) | 1.16 | (0.48) | 1.10 | (0.32) |
| Hispanic | 0.70* | (-2.24) | 0.71* | (-2.20) | 0.70* | (-2.20) | 0.71* | (-2.20) |
| Black | 0.83 | (-1.46) | 0.79 | (-1.82) | 0.77* | (-2.08) | 0.78* | (-1.97) |
| Asian | 0.35*** | (-3.42) | 0.36*** | (-3.30) | 0.34*** | (-3.36) | 0.36** | (-3.29) |
| Other Race | 0.60 | (-1.85) | 0.62 | (-1.76) | 0.57* | (-1.99) | 0.62 | (-1.76) |
| Female | 0.84 | (-1.73) | 0.89 | (-1.12) | 0.91 | (-0.87) | 0.91 | (-0.84) |
| Household Income | 1.06 | (1.05) | 1.05 | (0.92) | 1.04 | (0.74) | 1.05 | (0.91) |
| Parental Education | 1.15* | (2.34) | 1.13* | (2.02) | 1.15* | (2.30) | 1.15* | (2.22) |
| Age | 0.90** | (-3.04) | 0.91** | (-3.01) | 0.91** | (-2.99) | 0.91** | (-2.98) |
| Asexual Attraction | 1.00 | (-0.02) | 1.01 | (0.05) | 1.00 | (-0.01) | 1.00 | (0.02) |
| Bisexual Attraction | 0.71 | (-1.80) | 0.72 | (-1.67) | 0.74 | (-1.48) | 0.73 | (-1.63) |
| Same Sex Only Attract. | 0.65 | (-1.13) | 0.67 | (-1.08) | 0.64 | (-1.15) | 0.67 | (-1.08) |
| Severe Child Adversity | 0.98** | (-3.21) | 0.98** | (-3.11) | 0.98** | (-2.90) | 0.98** | (-3.12) |
| Illness |  |  | 0.79*** | (-4.84) | 0.80* | (-2.04) | 0.79** | (-3.13) |
| **Interaction effects** |  |  |  |  |  |  |  |  |
| Illness*Immigrant |  |  |  |  | 0.66 | (-1.44) |  |  |
| Illness*Hispanic |  |  |  |  | 1.03 | (0.24) |  |  |
| Illness*Black |  |  |  |  | 1.10 | (0.85) | 1.07 | (0.61) |
| Illness*Asian |  |  |  |  | 1.17 | (0.56) |  |  |
| Illness*Other Race |  |  |  |  | 1.26 | (0.97) |  |  |
| Illness*Female |  |  |  |  | 0.95 | (-0.52) | 0.96 | (-0.43) |
| Illness*Household Income |  |  |  |  | 1.03 | (0.58) |  |  |
| Illness*Parental Education |  |  |  |  | 0.92 | (-1.41) | 0.94 | (-1.23) |
| Illness*Asexual Attract. |  |  |  |  | 1.02 | (0.12) |  |  |
| Illness*Bisexual Attract. |  |  |  |  | 0.95 | (-0.26) |  |  |
| Illness*Same Sex Only Attract. |  |  |  |  | 1.22 | (0.62) |  |  |
| Illness*Severe Child Adversity |  |  |  |  | 1.00 | (-0.31) |  |  |
| Intercept | 18.50*** | (4.84) | 17.52*** | (4.78) | 17.39*** | (4.76) | 17.14*** | (4.74) |
| Random intercept variance | 8.58*** | (7.03) | 7.47*** | (6.83) | 7.48*** | (6.80) | 7.38*** | (6.79) |
| N | 9611 | | 9610 | | 9610 | | 9610 | |
| * p<0.05; ** p<0.01; *** p<0.001 | |  |  |  |  |  |  |  |

**Table S5b.** Main analysis using complete case analysis (rather than multiple imputation); Monogamous sex v. risky nonmonogamous sex comparison

|  | **Model 1** | | **Model 2** | | **Model 3** | | **Model 4** | |
| --- | --- | --- | --- | --- | --- | --- | --- | --- |
| **Main effects** | **OR** | **t** | **OR** | **t** | **OR** | **t** | **OR** | **t** |
| Immigrant | 1.34 | (1.12) | 1.26 | (0.89) | 1.36 | (1.14) | 1.26 | (0.89) |
| Hispanic | 0.88 | (-0.89) | 0.89 | (-0.84) | 0.86 | (-1.04) | 0.89 | (-0.84) |
| Black | 0.68** | (-3.27) | 0.65*** | (-3.74) | 0.62*** | (-4.03) | 0.63*** | (-3.91) |
| Asian | 0.64 | (-1.81) | 0.67 | (-1.61) | 0.62 | (-1.85) | 0.68 | (-1.57) |
| Other Race | 0.58* | (-2.20) | 0.59* | (-2.10) | 0.56* | (-2.23) | 0.60* | (-2.06) |
| Female | 1.36** | (3.25) | 1.46*** | (4.05) | 1.53*** | (4.39) | 1.53*** | (4.41) |
| Household Income | 0.99 | (-0.10) | 0.99 | (-0.27) | 0.98 | (-0.39) | 0.99 | (-0.27) |
| Parental Education | 1.21*** | (3.37) | 1.18** | (2.94) | 1.20** | (3.16) | 1.19** | (3.11) |
| Age | 0.96 | (-1.36) | 0.96 | (-1.33) | 0.96 | (-1.31) | 0.96 | (-1.29) |
| Asexual Attraction | 2.25*** | (5.33) | 2.29*** | (5.45) | 2.27*** | (5.27) | 2.28*** | (5.42) |
| Bisexual Attraction | 0.65* | (-2.42) | 0.67* | (-2.24) | 0.70 | (-1.93) | 0.68* | (-2.18) |
| Same Sex Only Attract. | 0.69 | (-1.11) | 0.71 | (-1.04) | 0.71 | (-1.04) | 0.72 | (-1.02) |
| Severe Child Adversity | 0.98*** | (-3.76) | 0.98*** | (-3.63) | 0.98*** | (-3.30) | 0.98*** | (-3.64) |
| Illness |  |  | 0.75*** | (-6.51) | 0.80* | (-2.16) | 0.77*** | (-3.75) |
| **Interaction effects** |  |  |  |  |  |  |  |  |
| Illness*Immigrant |  |  |  |  | 0.74 | (-1.32) |  |  |
| Illness*Hispanic |  |  |  |  | 1.14 | (1.00) |  |  |
| Illness*Black |  |  |  |  | 1.24* | (2.03) | 1.18 | (1.75) |
| Illness*Asian |  |  |  |  | 1.30 | (1.18) |  |  |
| Illness*Other Race |  |  |  |  | 1.19 | (0.81) |  |  |
| Illness*Female |  |  |  |  | 0.84* | (-1.98) | 0.85 | (-1.89) |
| Illness*Household Income |  |  |  |  | 1.00 | (0.06) |  |  |
| Illness*Parental Education |  |  |  |  | 0.93 | (-1.38) | 0.94 | (-1.47) |
| Illness*Asexual Attract. |  |  |  |  | 1.00 | (-0.02) |  |  |
| Illness*Bisexual Attract. |  |  |  |  | 0.93 | (-0.44) |  |  |
| Illness*Same Sex Only Attract. |  |  |  |  | 1.06 | (0.20) |  |  |
| Illness*Severe Child Adversity |  |  |  |  | 0.99 | (-1.15) |  |  |
| Intercept | 10.06*** | (4.15) | 9.59*** | (4.09) | 9.53*** | (4.08) | 9.42*** | (4.06) |
| Random intercept variance | 8.58*** | (7.03) | 7.47*** | (6.83) | 7.48*** | (6.80) | 7.38*** | (6.79) |
| N | 9611 | | 9610 | | 9610 | | 9610 | |
| * p<0.05; ** p<0.01; *** p<0.001 | |  |  |  |  |  |  |  |

**Table S5c.** Main analysis using complete case analysis (rather than multiple imputation); Abstinence v. risky nonmonogamous sex comparison

|  | **Model 1** | | **Model 2** | | **Model 3** | | **Model 4** | |
| --- | --- | --- | --- | --- | --- | --- | --- | --- |
| **Main effects** | **OR** | **t** | **OR** | **t** | **OR** | **t** | **OR** | **t** |
| Immigrant | 2.93*** | (4.32) | 2.68*** | (3.96) | 2.83*** | (4.04) | 2.65*** | (3.91) |
| Hispanic | 0.82 | (-1.47) | 0.83 | (-1.36) | 0.82 | (-1.44) | 0.83 | (-1.36) |
| Black | 0.36*** | (-8.73) | 0.34*** | (-9.36) | 0.33*** | (-9.50) | 0.33*** | (-9.45) |
| Asian | 0.76 | (-1.17) | 0.82 | (-0.86) | 0.77 | (-1.06) | 0.83 | (-0.80) |
| Other Race | 0.53** | (-2.69) | 0.55* | (-2.55) | 0.52** | (-2.69) | 0.55* | (-2.49) |
| Female | 0.98 | (-0.24) | 1.08 | (0.82) | 1.13 | (1.26) | 1.13 | (1.30) |
| Household Income | 1.14* | (2.54) | 1.12* | (2.29) | 1.11* | (2.08) | 1.12* | (2.29) |
| Parental Education | 1.33*** | (5.19) | 1.28*** | (4.61) | 1.30*** | (4.75) | 1.29*** | (4.69) |
| Age | 0.54*** | (-20.70) | 0.54*** | (-20.75) | 0.54*** | (-20.69) | 0.54*** | (-20.69) |
| Asexual Attraction | 3.34*** | (8.15) | 3.42*** | (8.32) | 3.37*** | (8.02) | 3.40*** | (8.29) |
| Bisexual Attraction | 0.54*** | (-3.60) | 0.56*** | (-3.35) | 0.58** | (-3.01) | 0.57** | (-3.25) |
| Same Sex Only Attract. | 1.01 | (0.03) | 1.04 | (0.13) | 1.04 | (0.11) | 1.05 | (0.15) |
| Severe Child Adversity | 0.95*** | (-8.89) | 0.95*** | (-8.75) | 0.95*** | (-8.27) | 0.95*** | (-8.77) |
| Illness |  |  | 0.68*** | (-9.10) | 0.77** | (-2.59) | 0.72*** | (-4.94) |
| **Interaction effects** |  |  |  |  |  |  |  |  |
| Illness*Immigrant |  |  |  |  | 0.77 | (-1.15) |  |  |
| Illness*Hispanic |  |  |  |  | 1.05 | (0.37) |  |  |
| Illness*Black |  |  |  |  | 1.34** | (2.77) | 1.29** | (2.72) |
| Illness*Asian |  |  |  |  | 1.19 | (0.82) |  |  |
| Illness*Other Race |  |  |  |  | 1.21 | (0.93) |  |  |
| Illness*Female |  |  |  |  | 0.76** | (-3.25) | 0.77** | (-3.17) |
| Illness*Household Income |  |  |  |  | 1.02 | (0.46) |  |  |
| Illness*Parental Education |  |  |  |  | 0.91 | (-1.87) | 0.93 | (-1.73) |
| Illness*Asexual Attract. |  |  |  |  | 1.07 | (0.50) |  |  |
| Illness*Bisexual Attract. |  |  |  |  | 0.99 | (-0.09) |  |  |
| Illness*Same Sex Only Attract. |  |  |  |  | 1.09 | (0.31) |  |  |
| Illness*Severe Child Adversity |  |  |  |  | 0.99 | (-1.65) |  |  |
| Intercept | 403722*** | (23.95) | 378012*** | (23.97) | 377437*** | (23.91) | 369103*** | (23.92) |
| Random intercept variance | 8.58*** | (7.03) | 7.47*** | (6.83) | 7.48*** | (6.80) | 7.38*** | (6.79) |
| N | 9611 | | 9610 | | 9610 | | 9610 | |
| * p<0.05; ** p<0.01; *** p<0.001 | |  |  |  |  |  |  |  |

**Table S6a.** Main text Table 2, Model 2 (protected nonmonogamous sex v. risky nonmonogamous sex comparison) modified by substituting the five individual (dichotomous) severe childhood adversities for the cumulative childhood adversity index; (MI, m=30)

|  | **Pooled** | | **Wave 1** | | **Wave 2** | |
| --- | --- | --- | --- | --- | --- | --- |
|  | **OR** | **t** | **OR** | **t** | **OR** | **t** |
| Immigrant | 0.976 | (-0.17) | 1.250 | (1.51) | 0.619* | (-2.22) |
| Hispanic | 0.773** | (-2.94) | 0.783** | (-2.62) | 0.799 | (-1.68) |
| Black | 0.862* | (-2.11) | 1.084 | (1.11) | 0.801* | (-2.17) |
| Asian | 0.512*** | (-4.42) | 0.526*** | (-3.99) | 0.506** | (-2.79) |
| Other Race | 0.824 | (-1.19) | 0.794 | (-1.35) | 0.955 | (-0.19) |
| Female | 0.868* | (-2.39) | 0.775*** | (-4.12) | 0.920 | (-0.91) |
| Household Income | 1.097** | (2.65) | 1.069 | (1.78) | 1.098 | (1.87) |
| Parental Education | 1.106** | (3.00) | 1.028 | (0.77) | 1.140* | (2.58) |
| Age | 0.976 | (-0.98) | 0.934*** | (-3.39) | 1.055 | (1.47) |
| asexual | 0.953 | (-0.44) | 0.853 | (-1.19) | 0.875 | (-0.91) |
| bisexual | 0.816 | (-1.82) | 0.889 | (-1.00) | 0.820 | (-1.11) |
| gay | 0.706 | (-1.48) | 0.831 | (-0.77) | 0.550 | (-1.44) |
| Sexual abuse | 0.837* | (-2.23) | 0.850 | (-1.72) | 0.984 | (-0.14) |
| Physical abuse | 1.005 | (0.05) | 0.977 | (-0.25) | 1.047 | (0.34) |
| Neglect | 0.788* | (-2.47) | 0.883 | (-1.28) | 0.749* | (-2.03) |
| Drugs/alcohol in home | 0.804** | (-2.72) | 0.901 | (-1.26) | 0.829 | (-1.54) |
| Parental incarceration | 0.911 | (-1.02) | 0.891 | (-1.31) | 1.019 | (0.13) |
| Intercept | 3.932** | (3.11) | 4.626*** | (4.39) | 0.588 | (-0.83) |
| Random intercept variance | 4.318*** | (9.58) |  |  |  |  |
| N | 41548 | | 20774 | | 20774 | |
| * p<0.05; ** p<0.01; *** p<0.001 | |  |  |  |  |  |

**Table S6b.** Main text Table 3, Model 2 (monogamous sex v. risky nonmonogamous sex comparison) modified by substituting the five individual (dichotomous) severe childhood adversities for the cumulative childhood adversity index; (MI, m=30)

|  | **Pooled** | | **Wave 1** | | **Wave 2** | |
| --- | --- | --- | --- | --- | --- | --- |
|  | **OR** | **t** | **OR** | **t** | **OR** | **t** |
| Immigrant | 1.196 | (1.46) | 1.532*** | (3.37) | 0.910 | (-0.57) |
| Hispanic | 0.904 | (-1.34) | 0.959 | (-0.50) | 0.906 | (-0.97) |
| Black | 0.627*** | (-7.23) | 0.824** | (-2.80) | 0.587*** | (-6.27) |
| Asian | 0.796 | (-1.88) | 0.802 | (-1.66) | 0.905 | (-0.57) |
| Other Race | 0.723* | (-2.18) | 0.724* | (-2.01) | 0.763 | (-1.30) |
| Female | 1.568*** | (8.58) | 1.323*** | (4.88) | 1.711*** | (7.18) |
| Household Income | 1.030 | (0.91) | 0.999 | (-0.04) | 1.030 | (0.69) |
| Parental Education | 1.139*** | (4.04) | 1.045 | (1.32) | 1.140** | (3.12) |
| Age | 1.070** | (3.27) | 0.924*** | (-4.26) | 1.084** | (2.95) |
| asexual | 2.450*** | (10.91) | 1.435** | (3.14) | 2.118*** | (7.37) |
| bisexual | 0.615*** | (-4.57) | 0.813 | (-1.86) | 0.546*** | (-3.86) |
| gay | 0.949 | (-0.26) | 0.706 | (-1.56) | 1.116 | (0.40) |
| Sexual abuse | 0.693*** | (-4.89) | 0.691*** | (-4.17) | 0.786* | (-2.16) |
| Physical abuse | 1.000 | (-0.00) | 1.032 | (0.36) | 0.975 | (-0.26) |
| Neglect | 0.875 | (-1.62) | 0.969 | (-0.35) | 0.845 | (-1.56) |
| Drugs/alcohol in home | 0.722*** | (-4.53) | 0.702*** | (-4.49) | 0.827* | (-2.01) |
| Parental incarceration | 0.881 | (-1.58) | 0.864 | (-1.73) | 0.922 | (-0.78) |
| Intercept | 1.397 | (0.91) | 6.594*** | (5.81) | 0.965 | (-0.08) |
| Random intercept variance | 4.318*** | (9.58) |  |  |  |  |
| N | 41548 | | 20774 | | 20774 | |
| * p<0.05; ** p<0.01; *** p<0.001 | |  |  |  |  |  |

**Table S6c.** Main text Table 4, Model 2 (abstinent v. risky nonmonogamous sex comparison) modified by substituting the five individual (dichotomous) severe childhood adversities for the cumulative childhood adversity index; (MI, m=30)

|  | **Pooled** | | | | | **Wave 1** | | | | **Wave 2** | |
| --- | --- | --- | --- | --- | --- | --- | --- | --- | --- | --- | --- |
|  | | **OR** | | **t** | | **OR** | | **t** | | **OR** | **t** |
| Immigrant | | | 2.221*** | (7.07) | | | 2.748*** | (8.82) | | 1.653** | (3.17) |
| Hispanic | | | 0.858* | (-2.04) | | | 0.905 | (-1.33) | | 0.856 | (-1.44) |
| Black | | | 0.350*** | (-16.56) | | | 0.406*** | (-14.12) | | 0.354*** | (-11.90) |
| Asian | | | 1.200 | (1.62) | | | 1.091 | (0.75) | | 1.450* | (2.23) |
| Other Race | | | 0.650** | (-3.13) | | | 0.669** | (-2.87) | | 0.677* | (-1.98) |
| Female | | | 1.446*** | (7.58) | | | 1.290*** | (4.90) | | 1.523*** | (6.15) |
| Household Income | | | 1.142*** | (4.31) | | | 1.112** | (3.22) | | 1.144** | (3.24) |
| Parental Education | | | 1.291*** | (8.31) | | | 1.217*** | (6.55) | | 1.279*** | (5.82) |
| Age | | | 0.612*** | (-26.50) | | | 0.544*** | (-35.82) | | 0.644*** | (-17.68) |
| Asexual Attraction | | | 3.535*** | (15.73) | | | 3.044*** | (10.85) | | 2.798*** | (10.16) |
| Bisexual Attraction | | | 0.611*** | (-5.03) | | | 0.735** | (-3.03) | | 0.571*** | (-3.78) |
| Lesbian/Gay Attraction | | | 0.955 | (-0.25) | | | 0.778 | (-1.25) | | 1.098 | (0.36) |
| Sexual abuse | | | 0.431*** | (-11.44) | | | 0.464*** | (-9.05) | | 0.464*** | (-7.28) |
| Physical abuse | | | 0.950 | (-0.69) | | | 0.969 | (-0.39) | | 0.932 | (-0.74) |
| Neglect | | | 0.654*** | (-5.35) | | | 0.688*** | (-4.67) | | 0.656*** | (-3.97) |
| Drugs/alcohol in home | | | 0.431*** | (-12.08) | | | 0.464*** | (-10.55) | | 0.467*** | (-8.35) |
| Parental incarceration | | | 0.623*** | (-6.16) | | | 0.625*** | (-5.95) | | 0.653*** | (-4.27) |
| Intercept | | | 50457.9*** | (32.90) | | | 160212.6*** | (40.87) | | 16401.8*** | (22.72) |
| Random intercept variance | | | 4.318*** | (9.58) | |  | | |  |  | |
| N | 41548 | | | | | 20774 | | | | 20774 | |
| * p<0.05; ** p<0.01; *** p<0.001 | | | | |  |  | | |  |  | |
